# The rich get richer: synaptic remodeling between climbing fibers and Purkinje cells in the developing cerebellum begins with positive feedback addition of synapses

**DOI:** 10.1101/627299

**Authors:** Alyssa Michelle Wilson, Richard Schalek, Adi Suissa-Peleg, Thouis Ray Jones, Seymour Knowles-Barley, Hanspeter Pfister, Jeff William Lichtman

## Abstract

During postnatal development, cerebellar climbing fibers strongly innervate a subset of their original Purkinje cell targets and eliminate their connections from the rest. In the adult, each climbing fiber innervates a small number of Purkinje cells and each Purkinje cell is innervated by a single climbing fiber. To get insight about the processes responsible for this remapping, we reconstructed serial electron microscopy datasets from mice during the first postnatal week. In contrast to adult connectivity, individual neonatal climbing fibers innervate many nearby Purkinje cells, and multiple climbing fibers innervate each Purkinje cell. Between postnatal days 3 and 7, Purkinje cells retract long dendrites and grow many proximal dendritic processes. On this changing landscape, individual climbing fibers selectively add many synapses to a subset of Purkinje cell targets in a positive-feedback manner, without pruning synapses from other Purkinje cells. The active zone sizes of synapses associated with powerful versus weak inputs are indistinguishable. These results show that changes in synapse number rather than synapse size are the predominant form of early developmental plasticity. Finally, although multiple climbing fibers innervate each Purkinje cell in early postnatal development, the number of climbing fibers and Purkinje cells in a local cerebellar region nearly match. Thus, initial over-innervation of Purkinje cells by climbing fibers is economical, in that the number of axons entering a region is enough to assure that each axon ends up with a postsynaptic target, and that none branched there in vain.

**HIGHLIGHTS:**

- Developing climbing fibers establish synapses on many neighboring Purkinje cells unlike the sparse pattern of innervation in later life
- Climbing fibers add many synapses onto a few of their Purkinje targets before the pruning stage in a rich-get-richer type process
- The synapse sizes of strengthened and weakened climbing fiber inputs are indistinguishable.
- Exuberant branching of climbing fiber axons in early postnatal life appears to be economical because the numbers of axons and Purkinje cells in a local region match, ensuring that each axon can establish a long-lasting connection there

**BLURB:** High-resolution serial electron microscopy reconstructions reveal that climbing fiber-Purkinje cell synaptic refinement in the developing cerebellum begins with significant synapse addition. Climbing fibers focus their synapses onto a smaller number of Purkinje cells by selectively adding synapses onto some target cells. All axons that project to a region in development play a role in the final connectivity.

## INTRODUCTION

In many vertebrates, neurons undergo extensive rewiring during postnatal development. Neurons remove synapses from some of their initial target cells, eventually achieving neural circuitry that is refined from what was initially an overconnected network. This process, known as synapse elimination, occurs in the central and peripheral nervous systems (CNS and PNS, respectively). One of the most striking examples of synapse elimination in the central nervous system occurs in the cerebellum, where connections between climbing fibers and Purkinje cells are refined. This phenomenon has been studied extensively in rodents, where shortly after birth, multiple climbing fibers innervate Purkinje cells in the cerebellar cortex (Crepel et al., 1976; Mariani and Changeux, 1981). Ultimately by the end of the third postnatal week in rodents only one climbing fiber innervates each Purkinje cell (for review see (Kano et al., 2018; Hashimoto and Kano, 2013)). The transition from multiple climbing fiber inputs to one parallels the most well known example of synapse elimination in the peripheral nervous system, which occurs between motor axons and muscle fibers at the neuromuscular junction. Perinatally, ~10 motor axons innervate each muscle fiber in a muscle (Tapia et al., 2012), but almost immediately after birth axons begin to remove their synapses from some muscle fibers. Live cell imaging *in vivo* shows that the remaining inputs increase their synaptic territory (synapse addition) through the takeover of sites occupied by other axons until only one axon innervates each muscle fiber (Walsh and Lichtman, 2003). This implies that at the neuromuscular junction, the addition of synapses is causally related to the vacation of sites occupied by the axons being pruned and supports the idea that this reorganization is based on a competition between axons vying to innervate the same postsynaptic cell. Time lapse imaging of developing neuromuscular junctions provided direct evidence that when one axon is eliminated experimentally it inevitably leads to compensatory synapse addition by a remaining input (Turney and Lichtman, 2012).

Cerebellar synapse elimination is more challenging to study because the cerebellar cortex is less accessible than the neuromuscular system, so that live imaging is difficult (Carrillo et al., 2013). In addition, climbing fibers and Purkinje cell geometries change considerably during early postnatal life when the connectivity is being refined (Chedotal and Sotelo, 1993; Ramón y Cajal, 1995 (1911)).

From an electrophysiological perspective it is clear that there are several stages of climbing fiber-Purkinje cell synaptic refinement during development. At around postnatal day 3 (P3), climbing fiber-Purkinje cell synapses become detectable in electrophysiological recordings (Mariani and Changeux, 1981). Several studies have estimated the number of climbing fibers innervating a single Purkinje cell to be typically fewer than 5 at this age, with all climbing fibers at a cell producing similar postsynaptic responses (Bosman et al., 2008; Mariani and Changeux, 1981; Scelfo and Strata, 2005). There is controversy over when this situation changes. Some work suggests that during the first postnatal week (in rodents) one of the recorded climbing fiber inputs to a Purkinje cell becomes more powerful than the others (Hashimoto and Kano, 2003; Bosman et al., 2008). However other researchers have found that this change does not occur until the second postnatal week (Scelfo and Strata, 2005) coincident with the initial loss of climbing fiber input. By the third week virtually every Purkinje cell is innervated by only one climbing fiber. The elimination process itself from P7 to beyond P10 also has been subdivided into three stages. First, between P7 and P9, some of the electrophysiologically weak climbing fiber inputs disappear. Second, at P9 to P10, the rise times associated with quanta of one of the remaining climbing fiber axons become longer (Hashimoto et al., 2009). Anatomical studies show that this phenomenon occurs as one climbing fiber begins its “climb” up the newly formed apical dendrite (Carrillo et al., 2013). Third, after P10, the few remaining climbing fibers that did not grow onto the apical dendrite progressively are lost and no longer elicit postsynaptic responses.

In this study, we aimed to resolve an important question about the rewiring of climbing fibers in development: are synapse strengthening and removal concurrent or sequential? Previous work has indicated that functional differentiation of climbing fibers occurs either in the first (Hashimoto and Kano, 2003; Bosman et al., 2008) or second (Scelfo and Strata, 2005) postnatal week, either before or during massive elimination of climbing fiber inputs, but cannot distinguish whether functional differentiation is a result of synaptic strengthening only or strengthening coupled with synapse removal. This is an important question because the answer provides insight into the underlying mechanism of developmental reorganization. For example, if synapse addition and elimination are occurring at the same time then, as at the neuromuscular junction, addition may be the consequence of loss; thus climbing fiber rearrangement could be a takeover-based competition. If however synapse addition is independent of synapse elimination, then the competitive mechanism described above for neuromuscular junction synapse elimination might be inadequate to explain the way this CNS reorganization takes place.

We also wanted to answer more specific questions about how changes in relative climbing fiber innervation strength are manifested. First, do climbing fibers alter their innervation strengths by adding and removing synapses, or by growing or shrinking individual synapses, or by a combination of these two phenomena? We are interested in this question because one of the limitations of the neuromuscular junction as a model of events in the CNS is that the neuromuscular junction looks different than typical synapses in the CNS. In the NMJ, where synapses grow through takeover, the synapse is a large, contiguous structure that has many presynaptic release sites and a quantal content of typically 100 (Slater, 2017). By contrast, CNS synapses are often single synaptic boutons with one or several release sites, each having quantal contents of ~1 (although some CNS synapses are substantially larger, e.g. calyx synapses with quantal contents of >300, (Iwasaki and Takahashi, 2001)). Thus climbing fiber synaptic strengthening could occur either by enlarging each synapse to have more release sites (by a small or large amount) or adding more boutons.

Furthermore, how does relative strengthening of climbing fiber synapses progress over time in the first postnatal week? And finally, how do climbing fibers parse out their synapses among many different Purkinje cells in the same local region of cerebellar cortex?

To explore these questions, we analyzed the connectivity of climbing fibers to Purkinje cells at P3 and P7 using serial-section scanning electron microscopy. We chose these time points because some evidence suggests that this is the stage when the first functional changes occur (Kano et al. 2018). Our results show that climbing fibers dramatically change their connectivity during this time, by adding synapses onto some target cells in a preferential manner. These results indicate that climbing fiber-Purkinje cell synaptic reorganization begins with axons choosing to establish additional synapses onto a small number of preferred partners among the cohort of Purkinje cells they innervate. In addition, we see no sign of any synapse loss. The connectivity we observe is well explained by a rich-get-richer process of synapse addition, also known as preferential attachment, in this case by climbing fibers onto their preferred targets. Finally, the connectomics approach also allowed us to learn about the population of axons that initially over-innervate the Purkinje cells in a region of cerebellar cortex. We find that the number of climbing fiber axons is approximately the same as the number of Purkinje cells locally (as is the case in the adult), implying that all the axons branching in an area are able to play a role in the final connectivity pattern.

## RESULTS

### The Datasets

These cerebellar datasets came from two developing mice, one at P3 and one at P7. Because we wished to compare them, for each we prepared a block from the medial cerebellar vermis and imaged lobule VIII, near the deepest part of the fissure (Figure 1a). We used the same region at each time point because the timeline for climbing fiber development is delayed for some lobules relative to others, and within each lobule timelines also shift with position (the outer region often develops earlier than the inner region (Armengol and Sotelo, 1991; Ramón y Cajal, 1995 (1911))). For both blocks, we imaged a column of cerebellar tissue reaching from the pia down to the white matter, in order to capture Purkinje cell dendrites and axons to the fullest extent possible. The axons were an easy way to identify Purkinje cells, since no other cell type consistently sends axons into the white matter. By focusing on the deepest part of the lobular fissure, we collected tissue closer to the proximal branch points of climbing fibers, which helped us reconstruct individual climbing fiber axons more fully. Each block was cut with an automatic tape-collecting ultramicrotome (ATUM) at 30-nm thickness in the sagittal plane (see Methods). A total of 1658 sections for the P3 block and 2514 sections from the P7 block were used for imaging in a scanning electron microscope. The sections were placed on wafers and images were acquired at 4 nm/px lateral resolution in a region measuring 190 μm x 120 μm, with the longer dimension being along the radial axis. The total acquired image volume was 1.1 x 10^6^ μm^3^ for P3 and 1.7 x 10^6^ μm^3^ for P7.

**Figure 1.**
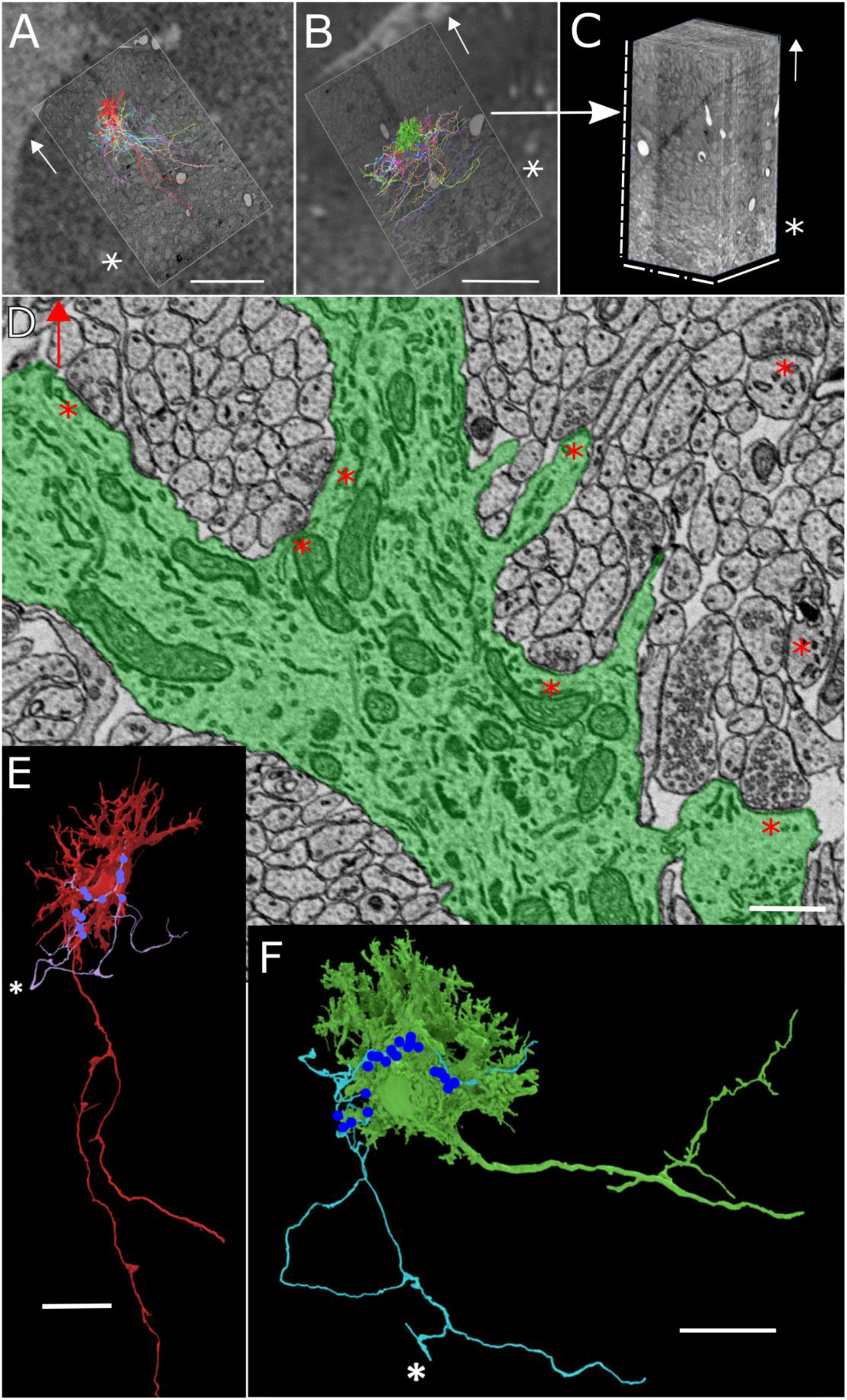
Reconstructions of Climbing Fiber Input to Purkinje Cells in P3 and P7 Mouse Cerebellar Cortex (a) A low-resolution electron micrograph of a single section from the P3 dataset. The subregion that was imaged at high resolution is superimposed onto the section. Also shown is the fully reconstructed Purkinje cell (red), along with all of the climbing fiber branches that innervate it. The arrow points in the direction of the pia and an asterisk indicates the location of the superficial white matter. (b) A low-resolution electron micrograph of a single section from the P7 dataset. The subregion imaged at high resolution is superimposed on the section. The fully reconstructed Purkinje cell (green) and all of its climbing fiber branch inputs are also shown. The arrow points in the direction of the pia and the asterisk indicates the location of the superficial white matter. The cortical layers for the P3 and P7 datasets are shown in Figure S1a. The other Purkinje cells reconstructed in both volumes are shown in Figures S1b and S1c. Closer views of the fully reconstructed Purkinje cells, showing their topography, are shown in Figures S1d and S1e. (c) The electron micrograph image volume collected in P7 cerebellum (b). The solid line denotes the direction through tissue sections. White features in the section images are blood vessels. The arrow points toward the pia, and the asterisk denotes the location of the superficial white matter. (d) A region from a high-resolution micrograph from one of the sections imaged at P7. Part of a dendrite from the fully reconstructed Purkinje cell has been manually traced (green). In this section multiple excitatory synapses are visible (asterisks). The arrow points in the direction of the pia. (e) The fully reconstructed Purkinje cell at P3, shown with its most powerful climbing fiber branch input (purple). Dots show the locations of synapses (13 in total) formed by the climbing fiber branch onto the Purkinje cell. The point at which the climbing fiber branch enters the volume is shown by an asterisk. (f) The fully reconstructed Purkinje cell at P7, shown with its most powerful climbing fiber branch input (blue). Dots show the locations of synapses (26 in total) formed by the climbing fiber branch onto the Purkinje cell. The point at which the branch enters the volume is shown by an asterisk. Scale bars: (a), (b) 100 μm; (c) 75 μm solid line, 120 μm dot-dashed line, 190 μm dashed line; (d) 0.5 μm; (e), (f) 15 μm.

### Cytoarchitecture of the Cerebellum at Two Developmental Ages

All layers of the cerebellum were visible in both the P3 and P7 datasets although the thicknesses of some layers differed between the two ages, as expected (Figure S1a). The molecular layer was substantially thicker at P7 than at P3, consistent with granule cell migration from external to internal granular layers (Sotelo, 2004). Additionally, the Purkinje cell layer was 2 to 3 cells deep at P3 and became a true monolayer at P7 (Figure S1b). The density of Purkinje cells in the Purkinje cell layer also decreased from P3 to P7 (0.008 cells/μm^2^ at P3 to 0.003 cells/μm^2^ at P7, possibly due to tissue growth). We also observed substantial changes in Purkinje cell geometry (see below).

### Changing Connectivity Patterns on Cerebellar Purkinje Cells between P3 and P7

In order to reveal how Purkinje cell innervation changed during the first postnatal week, we sampled climbing-fiber-Purkinje cell connectivity in both volumes. We began this sampling by reconstructing all synaptic input to one Purkinje cell at each age (using a combination of manual and automatic segmentation--see Methods). This task was challenging because the dendritic arbors of the immature Purkinje cells were quite complex. At P3 (Figure S1c, left panel), dendrites ran in many different directions (in dramatic contrast to the monoplanar geometry of adult Purkinje cell dendrites) and made frequent side branches. Because of this geometry, the dendritic arbor ramified over a large portion of the image volume, meaning that identifying synapses onto the Purkinje cells required scrutiny of nearly the entire volume. We found a total of 464 synapses targeting the P3 cell. At P7 (Figure S1c, right panel), although the Purkinje cell had about 3 times fewer dendrites, both dendrites and soma produced a large number of spine-like processes (as reviewed in (Sotelo, 2004)), many of which were innervated. (Interestingly, a number of these processes ran parallel with nearby parallel fibers--data not shown.) Finding synapses on the P7 cell thus involved tracing out thousands of processes. In total, we found 1048 synapses on the P7 cell.

To identify the climbing fiber inputs to these Purkinje cells, we classified the presynaptic partner for every synapse based on the input axon’s morphology and its synaptic connections to other target cells. Using this procedure, we found that 352 axon branches formed all the synapses onto the fully reconstructed Purkinje cell in the P3 data, and 693 axon branches formed the synapses onto the fully reconstructed Purkinje cell at P7 (see Table 1). At both ages it was possible to divide the axons into three general categories: inhibitory cell axons, granule cell axons (either the initial radial portion or parallel fibers), and non-granule-cell excitatory axons. Non-granule-cell excitatory axons may be either climbing fibers (olivocerebellar axons) or mossy fibers (axons from a number of sources). We wanted to distinguish between the mossy and climbing fiber input, because the main reorganization concerns selection of one climbing fiber input and elimination of the rest, whereas the mossy input is minor during development and then disappears entirely (Mason and Gregory, 1984; Kalinovsky et al., 2011). During the first postnatal week it was difficult to distinguish these two axon types based on their morphology (although they become easily distinguishable within a few days after this period--see Mason and Gregory, 1984). We thus attempted to separate them by their synaptic connectivity. We analyzed the full synaptic connectivity of the excitatory non-granule axon branches, and found that they formed two groups, one that formed a majority of their synapses onto Purkinje cells and one that formed a majority of their synapses onto granule cells. To determine whether the second group were mossy fibers, which target mainly granule cells, we compared the connectivity of the Purkinje cell-innervating axons with that of a sample of putative mossy fibers (that we identified by virtue of the fact that they innervated granule cell dendrites deep in the internal granular layer, a pattern typical of mossy fibers). We found by several different clustering methods (see Methods, Figure S2) that the non-granule excitatory axon branches that innervated our fully reconstructed Purkinje cells fell into two groups: those that resembled mossy fibers and those that resembled climbing fibers. Through this analysis we identified 55 climbing fiber axon branches from the 84 non-granule excitatory inputs to the fully reconstructed Purkinje cell at P3, and 49 climbing fiber branches from the 64 non-granule excitatory inputs to the Purkinje cell at P7 (see examples in Figures 1e and 1f).

**Table 1.** Summary of Synaptic Inputs to Fully Reconstructed Purkinje Cells at P3 and P7

|  | # Synapses from inputs, total | # Synapses, CFs (% of all synapses from all inputs) | # Inputs, total | # Climbing fiber branches (% of total number of inputs) | # Mossy fiber branches (%) | # Non-granule excitatory branches forming < 5 synapses (%) | # Parallel fibers (%) | # Granule cell axon branches (%) | # Inhibitory axon branches (%) |
| --- | --- | --- | --- | --- | --- | --- | --- | --- | --- |
| P3 | 464 | 127 (27.3%) | 352 | 55 (15.6%) | 12 (3.4%) | 17 (4.8%) | 259 (73.6%) | 1 (0.3%) | 8 (2.3%) |
| P7 | 1048 | 298 (28.4%) | 693 | 49 (7.1%) | 13 (1.9%) | 2 (0.3%) | 619 (89.3%) | 4 (0.6%) | 6 (0.9%) |

We examined the synaptic connections of all climbing fiber branches innervating the fully reconstructed Purkinje cells at P3 and P7. Despite the fact that a slightly smaller number of climbing fiber axon branches innervated the P7 Purkinje cell compared with the P3 cell (49 vs. 55; probably insignificant—see below), there was a greater-than-twofold increase in the total number of synapses formed by those climbing fibers (127 at P3 to 298 at P7). This increase in synapse number was a general feature of individual climbing fiber axon branches: the number of synapses formed by each climbing fiber branch at P3 was significantly smaller than the number of synapses per branch at P7 (Figures 2a and 2b).

**Figure 2.**
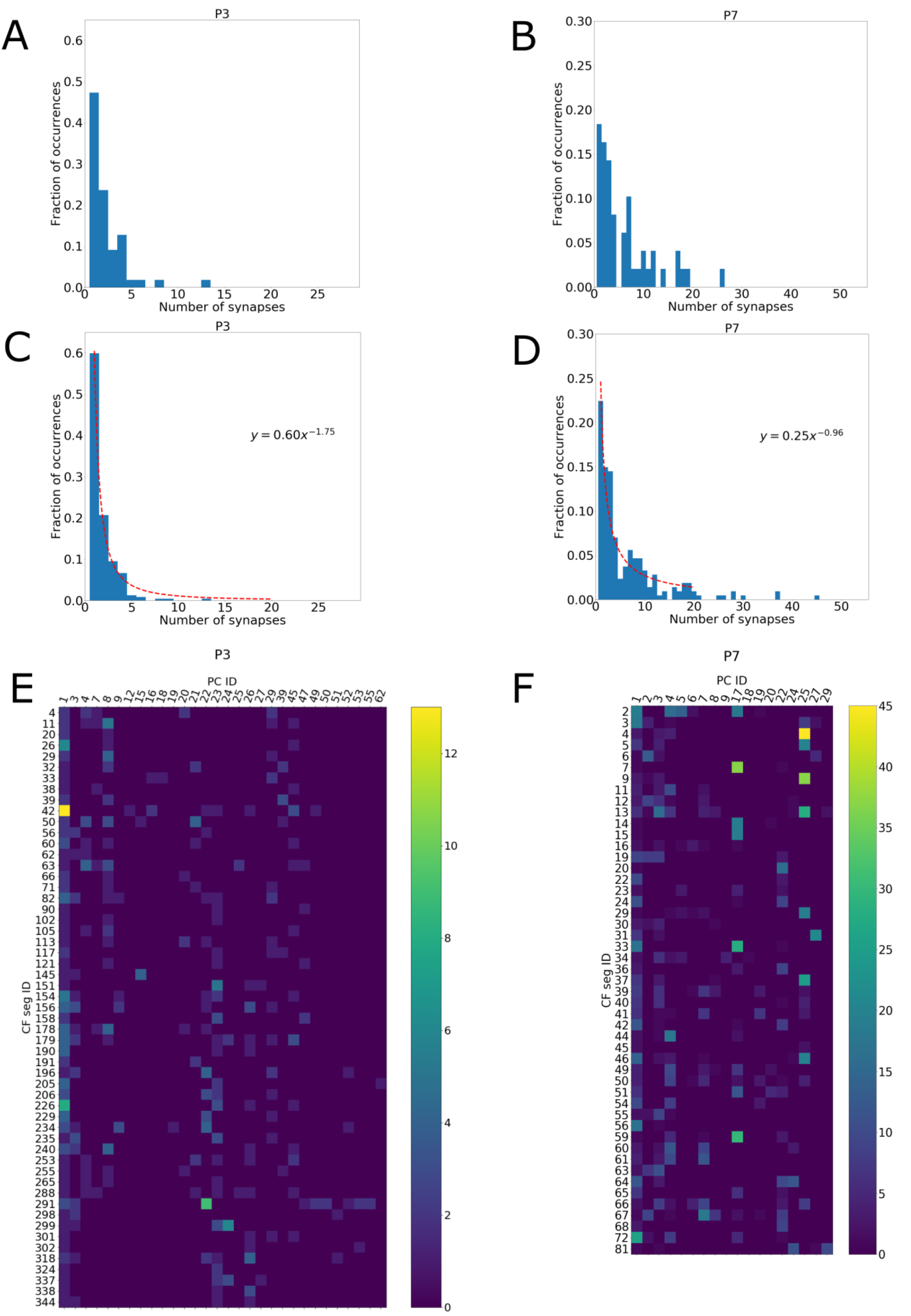
A Small Proportion of Climbing Fiber Branch-Purkinje Cell Connections Consist of Disproportionately Large Numbers of Synapses (a) Histogram showing the number of synapses formed by all climbing fiber inputs to the fully reconstructed Purkinje cell at P3. The 55 climbing fiber branch inputs formed a total of 127 synapses onto the fully reconstructed Purkinje cell. The mean +/− standard deviation for this tailed distribution was 2.3 +/− 2.1 synapses per connection, and the median was 2 synapses per connection. (b) Histogram showing the number of synapses formed by all climbing fiber branch inputs to the fully reconstructed Purkinje cell at P7. The 49 climbing fiber branch inputs formed a total of 298 synapses onto the fully reconstructed Purkinje cell. The mean +/− standard deviation for this tailed distribution was 6.08 +/− 5.7 synapses per connection, and the median was 4 synapses per connection. (c) Histogram showing the number of synapses formed between all climbing fiber branch-Purkinje cell pairs reconstructed in the P3 image volume. The 55 climbing fiber branches that innervated the fully reconstructed Purkinje cell innervated 63% (30/48) of all Purkinje cells in the volume, forming 242 climbing fiber-Purkinje cell pairs and a total of 435 synapses with those targets. The mean +/− standard deviation of this tailed distribution was 1.8 +/− 1.4 synapses per connection, and the median was 1 synapse per connection. (d) Histogram showing the number of synapses formed between all climbing fiber branch-Purkinje cell pairs reconstructed in the P7 image volume. The 49 climbing fibers that innervated the fully reconstructed Purkinje cell at P7 innervated a total of 60% (18/30) Purkinje cells in the volume, forming 214 climbing fiber-Purkinje cell pairs, and 1355 synapses with those targets. The mean +/− standard deviation for this distribution was 6.3 +/− 7.1 synapses per connection, and the median was 3 synapses per connection. Monte Carlo simulations show that these distributions cannot result from climbing fiber branches innervating Purkinje cell targets via a uniform probability distribution (see Figure S5). (e) Connectivity matrices for all reconstructed climbing fiber branches onto all Purkinje cell targets at P3. Each row represents a single climbing fiber branch and each column represents a single Purkinje cell. The value in the *i*th row and *j*th column is the number of synapses formed by climbing fiber branch *i* onto Purkinje cell *j*. (f) Connectivity matrices for all reconstructed climbing fiber branches onto all Purkinje cell targets at P7. The value in the *i*th row and *j*th column is the number of synapses formed by climbing fiber branch *i* onto Purkinje cell *j*.

The differing behavior we observed for climbing fiber axon branches at P3 and P7 was confirmed when we expanded our connectivity sample, by including all the synapses formed by these axon branches onto other Purkinje cells in the volume (Figures 2c-f). The 55 climbing fiber branches in the P3 dataset innervated 29 of the 47 other Purkinje cells in the volume (in total, 30/48 or 63% of Purkinje cells) and allowed analysis of 187 additional climbing fiber-Purkinje cell connections (consisting of 308 additional synapses). The 49 climbing fiber branches in the P7 dataset innervated 17 of the 29 other Purkinje cells in the volume (a total of 18/30 or 60% of Purkinje cells), contributing 165 additional climbing fiber-Purkinje cell connections and 1057 additional synapses. For both ages, the extended data corroborated the analysis described above (Figures 2a-d): axons were establishing greater numbers of synapses at P7 than at P3.

Furthermore, at both ages these axons distributed their synapses amongst Purkinje cells in an inhomogeneous way, in which most climbing fiber-Purkinje cell connections consisted of a few synapses, and a small number of climbing fiber-Purkinje cell connections consisted of many synapses (Figure 2). We considered two different ways that climbing fiber branches could generate the synapse distributions we observed. One possibility is that each individual climbing fiber branch forms an inhomogeneous distribution of synapses, establishing many synapses with a few Purkinje cell targets and a few synapses with the rest. Alternatively, some climbing fiber branches might establish few synapses with every one of their targets, whereas other climbing fiber branches establish many synapses with each of their targets. To decide between these alternatives, we inspected the number of synapses formed by each climbing fiber branch onto each of its Purkinje cell targets (Figures 3a, S3, and S4). At both P3 and P7 we found that all climbing fiber branches formed only one or two synapses onto most of their Purkinje targets, and at both ages some axons also innervated a few cells with larger numbers of synapses. In neither dataset did we observe a climbing fiber that formed only large numbers of synapses onto its Purkinje cell targets. This data argues that developing climbing fibers tend to behave similarly to each other, innervating a large number of Purkinje cells with a few synapses and innervating a few Purkinje cells with many synapses. Furthermore, the preferences we observed in these distributions could not have occurred if climbing fibers distributed their synapses amongst their Purkinje cell targets uniformly (Methods and Figure S5; p < 0.00001 for both P3 and P7).

**Figure 3:**
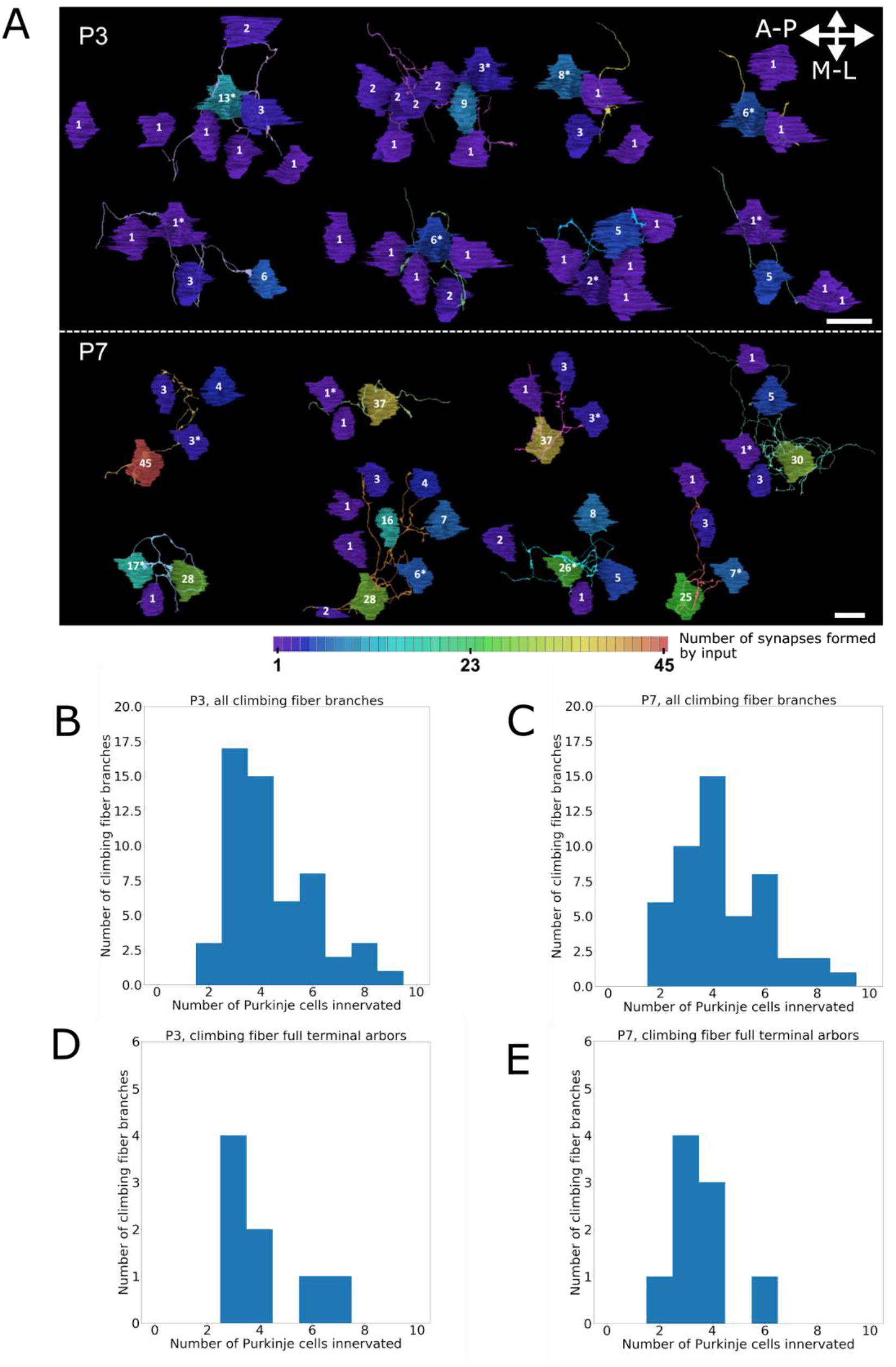
Synaptic Divergence Patterns for Climbing Fiber Branches at P3 and P7 (a) Connectivity onto Purkinje cell targets for the eight climbing fiber branches that form the largest numbers of synapses onto a single Purkinje cell at P3 (top panel) and P7 (bottom panel). Each climbing fiber branch is shown from a radial view (looking down at the white matter from the pia). The anteroposterior axis (which falls in the sagittal plane) and mediolateral axis are shown with arrows. Each climbing fiber branch is shown with the Purkinje cells it innervates (only somas are shown for clarity), and the number of synapses it forms onto each cell is shown on the soma. The color of a Purkinje cell also indicates the number of synapses formed onto it, with colors ranging from dark purple for 1 synapse to red for the maximum observed number of synapses (45). See also Figures S3, S4, and S7. Climbing fiber branch IDs at P3, in order of decreasing maximum number of synapses, are 42, 291, 226, 26 (left to right, top row), 299, 154, 11, 151 (left to right, bottom row). Climbing fiber branch IDs at P7, in order of decreasing maximum number of synapses, are 4, 7, 9, 59 (left to right, top row), 33, 13, 72, 37 (left to right, bottom row). (b) Histogram showing the number of Purkinje cells innervated by each climbing fiber branch reconstructed at P3 (55 climbing fiber branches; mean +/− standard deviation = 4.4 +/− 1.6) (c) Histogram showing the number of Purkinje cells innervated by each climbing fiber branch reconstructed at P7 (49 branches; mean +/− standard deviation = 4.4 +/− 1.7). (d) Histogram showing the number of Purkinje cells innervated by only climbing fiber branches whose full terminal arbors are contained within the P3 image volume (8/55 branches; mean +/− standard deviation = 4.1 +/− 1.5). (e) Histogram showing the number of Purkinje cells innervated by only climbing fiber branches whose full terminal arbors are contained within the P7 image volume (9/49 branches; mean +/− standard deviation = 3.6 +/− 1.6). Scale bars: (a) 15 μm.

This tendency for axons to distribute a large number of synapses on a small number of target cells was more extreme at P7 than P3. At P3, the largest number of synapses we observed a climbing fiber branch to form was 13, but at P7 the largest number of synapses was 45. This age difference implies that axons are focusing progressively more synapses on the target cells they prefer.

We next investigated the spatial organization of the uneven distribution of synapses formed by individual climbing fiber branches. In general, climbing fiber branches at P7 formed a higher density of synapses than at P3 (Figure S6a; Wilcoxon rank sum test, p = 3×10^−11^). At both ages climbing fibers formed synapses onto Purkinje cells somewhat sparsely and at regular intervals as they ramified through the Purkinje cell layer (P3: https://github.com/amwilson149/baby-andross/blob/master/Neuroglancer_links/190502_p3_CFs_and_syns_MFs_and_syns_PC1_and_all_PC_somas.txt; P7: https://github.com/amwilson149/baby-andross/blob/master/Neuroglancer_links/190502_p7_CFs_and_syns_MFs_and_syns_PC1_and_all_PC_somas.txt [links will be updated]; see Figure S7a for an example from P7). The one exception to this trend was that some climbing fiber branches with strong Purkinje cell preferences formed higher-density clusters in localized regions (onto the preferred Purkinje cells; Figure S7a). The Purkinje cells that received large numbers of synapses in these clusters were located at various positions relative to the climbing fiber branch itself. Specifically, the preferred cells of a climbing fiber branch did not necessarily overlap maximally with that branch’s arbor. In fact, at both ages, climbing fiber branches strongly innervated Purkinje cells in a salt and pepper pattern, often making no or very few synapses on an immediate neighbor of a strongly innervated cell (e.g. Figure S7b). The discontinuous innervation patterns we observe suggest that the number of synapses formed by a climbing fiber onto a Purkinje cell is based on a form of selectivity that is not related to location of the individual Purkinje cell targets.

At both P3 and P7, the distributions of synapses established by individual climbing fibers onto Purkinje cells appear to obey a power law. Power laws arise in many systems that undergo growth (see Newman, 2005 for a review). The power law distribution (*P*(*k*) = *Ck*^−α^, where *C* and *α* are constants) determines the proportion of objects *P*(*k*) in a system (in this case, synaptically connected climbing fiber-Purkinje cell pairs) that have a particular number *k* of some quantity (in this case, synapses from the climbing fiber branch onto the Purkinje cell). Both synapse distributions were well fit by power law functions (Figures 2c and 2d) with *α* = *1.75* for P3 and *α* = *0.96* for P7 (the lower value at P7 corresponds to a slower drop-off in the number of climbing fiber-Purkinje cell pairs that share increasing numbers of synapses). These power law distributions could arise from a particular type of growth process known as preferential attachment (Barabási et al., 1999). In the climbing fiber-Purkinje cell system, preferential attachment would mean that the tendency of climbing fibers to add synapses increases as the number of synapses they already share with a Purkinje cell target grows larger.

Although there is evidence of massive synaptic pruning in the second postnatal week (measured as a decrease in the number of climbing fiber inputs to a Purkinje cell—see (Hashimoto and Kano, 2003; Bosman et al., 2008; Scelfo and Strata, 2005)), we were interested to learn whether our detailed electron microscopy data showed evidence of climbing fibers removing synapses from Purkinje cells while they added synapses onto others during the first postnatal week. Because we saw many single- and few-synapse inputs, synapse removal would likely result in some decrease in the number of Purkinje cells innervated by a climbing fiber branch. Therefore, at P3 and P7, we compared the number of Purkinje cells innervated by each climbing fiber branch (Figures 3b and 3c). The distributions at P3 and P7 were not significantly different (Wilcoxon rank-sum test, p = 0.99). However, both of these histograms included information from climbing fiber branches that had terminal arbors that were only partially contained inside the volume (that is, whose most distal branches ramified outside the volume boundary at some point), and furthermore, the P3 and P7 image volumes were not identical in size. Both of these factors could obscure an actual change in the divergence between P3 and P7, especially if that change is subtle. To make a more reliable measurement of the number of Purkinje cells a climbing fiber innervates (i.e. its divergence), we removed from our analysis all climbing fiber branches whose terminal arbors left the volume (leaving 8/55 climbing fiber segments at P3 and 9/49 at P7; Figures 3d and 3e). The divergences we measured for these branches with complete terminal arbors also did not differ significantly between P3 and P7 (Wilcoxon rank-sum test, p = 0.58), arguing that climbing fiber branches remain connected to roughly the same number of Purkinje cells between P3 and P7 and were hence not undergoing net pruning over these four days. For this reason, we suspect that the smaller number of climbing fiber branches we observed to innervate the fully reconstructed P7 cell compared with the P3 cell (see above) is likely due to random variation in climbing fiber innervation and in the different sizes of the two datasets, rather than a systematic change in connectivity between these two ages.

To summarize, the main difference between the P3 and P7 datasets is that individual climbing fibers form significantly more synapses onto their Purkinje cell targets at P7 than at P3. Notably, the synapse addition process that occurs between these two ages appears to be preferential, because climbing fibers elaborate many synapses onto a small number of their Purkinje cell targets.

### Strong and Weak Climbing Fiber Inputs to Purkinje Cells Have the Same Size Synapses

As shown by several studies, the extent of the postsynaptic density (PSD) can serve as a proxy for synaptic strength (Bailey and Chen, 1983; Bailey and Chen, 1988a, b; Bartol et al., 2015). We therefore measured the PSD areas for all identified climbing fiber-Purkinje cell synapses at each age (see Methods, Figures 4a-c). The PSD area distributions were significantly different between P3 and P7 (Figure 4d, p ≈ 0, Wilcoxon rank sum test). In particular, although the range of PSD areas overlapped, in the younger mouse the median PSD area was 1.5 times larger, and the variance was 4.2 times larger than at P7 (median of 0.22 μm^2^ at P3 vs. 0.088 μm^2^ at P7; variance of 0.14 μm^2^ at P3 vs. 0.019 μm^2^ at P7). One interpretation of this difference is that as the nervous system matures, synapse size becomes more consistent. To learn whether climbing fiber branches that form the smallest numbers of synapses onto a Purkinje cell (and may be more likely to be pruned away) have PSD areas that differ significantly from climbing fiber branches that form large numbers of synapses onto a Purkinje cell (and hence may be more likely to remain), we measured the correlation between the PSD area of a synapse and the total number of synapses shared by its pre- and postsynaptic partners. We found that synaptic area was not correlated with the number of synapses formed between a climbing fiber and Purkinje cell (Figures 4e-h; p = 0.4 at P3 and p = 0.7 at P7 for permutation tests performed with 100,000 permutations each). Taken together, these results show that individual climbing fiber-to-Purkinje cell synapses are indistinguishable from a PSD perspective and suggest that the primary means by which a climbing fiber changes its efficacy is by altering synapse number.

**Figure 4.**
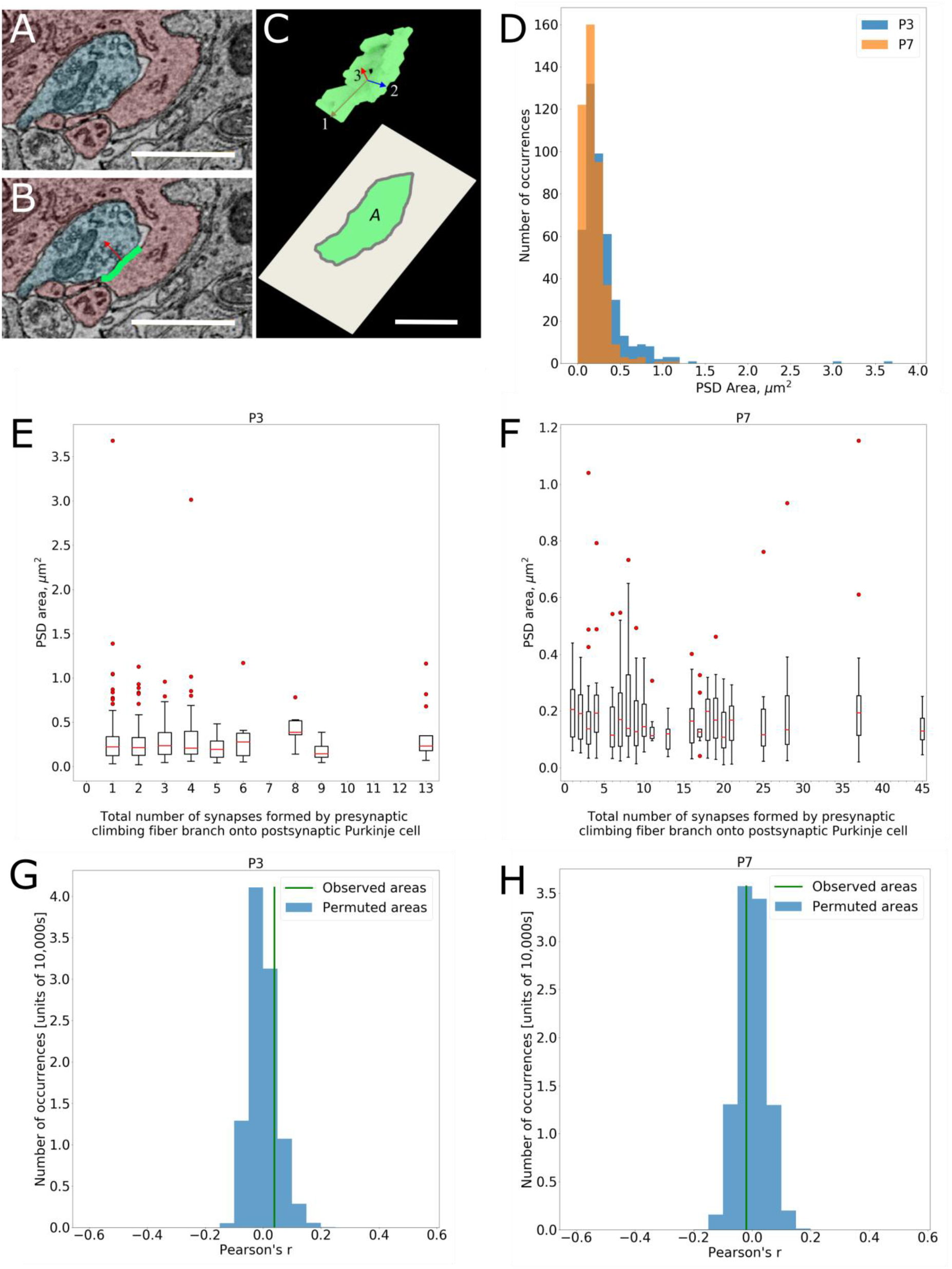
PSD Areas Become More Uniform During the First Postnatal Week and are Uncorrelated to Climbing Fiber-Purkinje Cell Connection Strength (a) Region of a high-resolution electron micrograph showing a synapse formed by a climbing fiber branch (blue) onto a Purkinje cell (red). (b) Same electron micrograph region as in (a), with a manual annotation of a PSD formed by the climbing fiber branch onto the Purkinje cell at the synapse shown. The red arrow points in the direction normal to the flattest plane of the PSD. (c) Top: a three-dimensional rendering of the PSD annotation for the synapse shown in (a) and (b). Three arrows show the first, second, and third components that result from applying principal component analysis to the PSD voxel list. The third component is used as the direction of the flattest plane of the PSD. A two-dimension projection of the PSD along that axis (bottom) is then used to estimate the PSD area, *A*, for the synapse. (d) Histograms of the PSD areas measured for all climbing fiber-Purkinje cell synapses identified at P3 (blue, 435 synapses) and P7 (orange, 1355 synapses). (e) Box plot summarizing the PSD area for each synapse vs. the total number of synapses formed between its presynaptic climbing fiber branch and postsynaptic Purkinje cell at P3. (f) Box plot summarizing the PSD area for each synapse vs. the total number of synapses formed between its presynaptic climbing fiber branch and postsynaptic Purkinje cell at P7. (g) Results of a permutation test comparing the correlation observed at P3 between PSD area and climbing fiber branch-Purkinje cell connection strength (green line, from (e)) against correlations measured when PSD areas are randomly shuffled (blue histogram). (h) Results of a permutation test comparing the correlation observed at P7 between PSD area and climbing fiber branch-Purkinje cell connection strength (green line, from (f)) against correlations measured when PSD areas are randomly shuffled (blue histogram). Scale bars: (a), (b) 1 μm, (c) 0.25 μm.

### Simulation: Synapse Addition via Preferential Attachment

To better understand the process by which the P3 connectivity pattern transformed into the pattern observed at P7, we constructed a time-stepping simulation to evolve the connectivity we observed at P3 to a state of connectivity similar to that at P7. The model (see Methods for details) allowed stochastic removal and addition of synapses by each climbing fiber branch with the Purkinje cells it was already connected to. Because we found that climbing fibers establish disproportionately large numbers of synapses onto subsets of their Purkinje targets (see above), the model also controlled the probability of choosing a particular target cell for synapse addition. This probability was proportional to the number of synapses currently shared by the climbing fiber and Purkinje cell target. The exponent of this proportionality, γ, was varied to reflect a range of behaviors, from preferential attachment (positive γ), to no dependence (γ = 0), to penalization of strong inputs (negative γ).

Varying (1) the relative rates of synapse addition and removal and (2) the value of γ allowed us to test a range of potential models for climbing fiber-Purkinje cell synapse rewiring. We considered a model to have achieved P7-like connectivity if the P3 connectivity matrix (Figure 2c) evolved into one whose size, overall synapse distribution, climbing-fiber-wise (rows) synapse distribution, and Purkinje-cell-wise (columns) synapse distribution became statistically indistinguishable from those of the observed P7 connectivity matrix (Figure 2d), based on Wilcoxon rank-sum tests at an alpha level of 0.05.

We went through this exercise to test in particular two different mechanisms that might produce the transformation between P3 and P7 connectivity. First, this transformation might occur through preferential attachment of climbing fibers to their target cells, in which strong connections preferentially get stronger (add more synapses). In the simulation, this behavior would be represented by a positive value of γ. Alternatively, the P7 result could potentially come about without preferential attachment (that is, with a non-positive γ), if there were both non-selective addition (γ = 0) and elimination of synapses. In this case, the number of synapses shared by any climbing fiber-Purkinje cell pair would do a random walk over time about its P3 value. As a result, some pairs would “drift” towards larger numbers of shared synapses and other pairs would drift towards smaller numbers, causing a spread in the histogram of synapse numbers, which qualitatively is what we observed at P7.

We found, however, that only preferential attachment processes (Newman, 2005; Barabási et al., 1999) could actually evolve P3 connectivity into P7-like connectivity (Figure 5a). Non-selective addition and removal of synapses always evolved the P3 distribution to one that was less like a power law and thus never converged toward the actual P7 distribution (Figure S8a-d). With preferential attachment, however, P3 connectivity evolved into a state statistically indistinguishable from P7 if the rate of synapse removal was zero or low (at least 2000 times smaller than the rate of addition), and if γ was roughly in the range of 0.5 to 1.7 (Figure 5a inset). These parameter values suggest that removal of synapses is not a significant component of synapse rewiring between P3 and P7 (consistent with our studies of axonal divergence; see above). These values also argue that preferential synapse addition by climbing fibers onto their Purkinje cell targets is required to produce the disproportionately large numbers of synapses we observed at P7.

**Figure 5.**
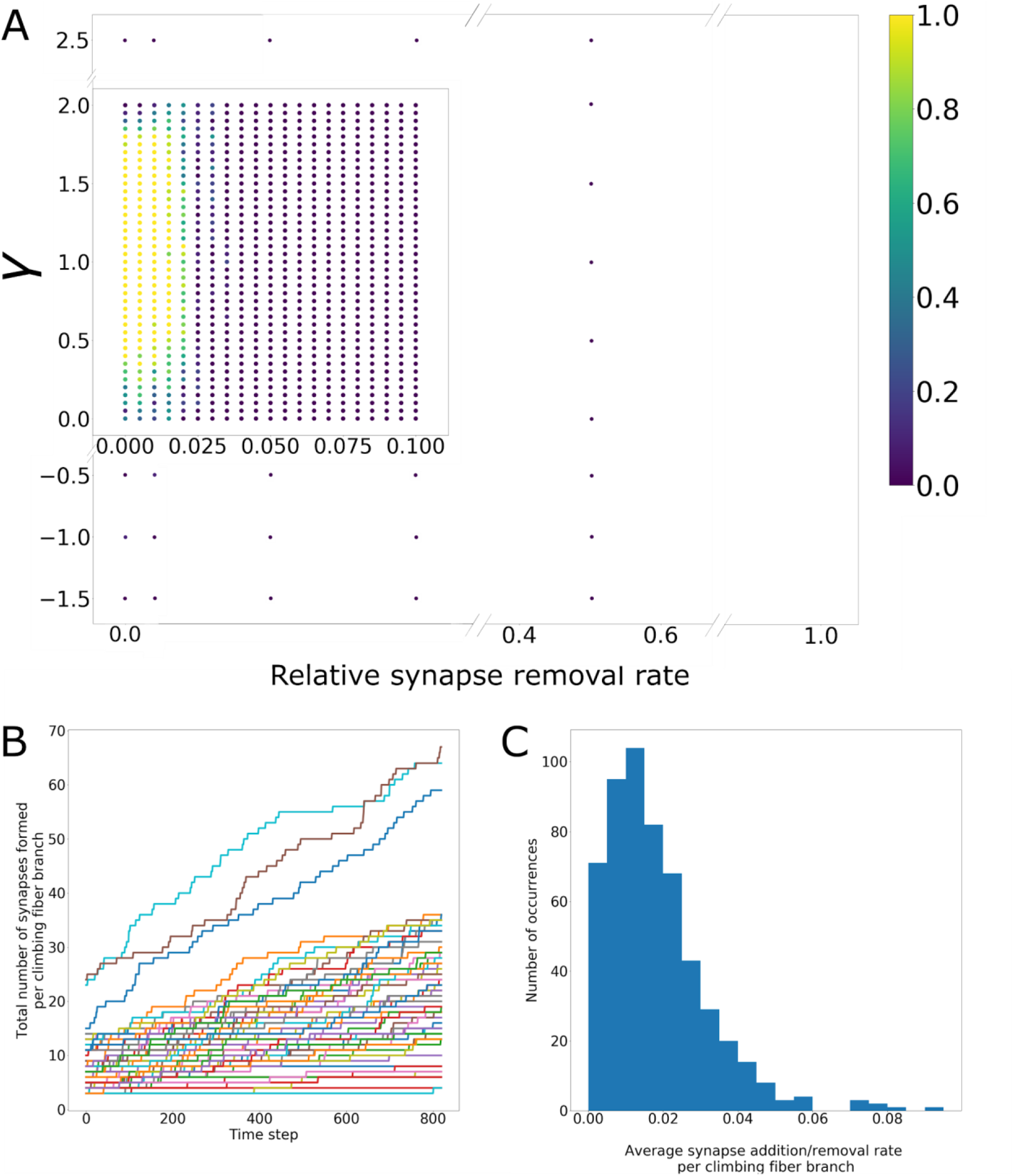
Preferential Addition of Synapses and Minimal Synapse Removal are Required to Evolve P3 Connectivity to P7 Connectivity (a) Plot showing the fraction of times that the simulation of stochastic synapse addition and removal was able to evolve the observed P3 connectivity matrix into one statistically indistinguishable from the observed P7 connectivity matrix. For each pair of parameter values, the fraction of convergent simulations was computed based on the results of 10 simulation runs. (b) Plot of the total number of synapses formed by each climbing fiber branch at each time step of the stochastic synapse addition and removal simulation, for a convergent parameter pair (*p_rem_* = 0.005, *ɣ* = 1.1). Time steps are shown up until the first time step where P7-like connectivity was achieved. For each climbing fiber branch, the average rate of synapse addition was computed from this data by taking the slope of the best-fit line. (c) The histogram of average synapse addition rates per climbing fiber branch, compiled from 100 runs of the simulation with the same convergent parameters as in (b) (*p_rem_* = 0.005, *ɣ* = 1.1). The median and maximum of this distribution are 0.015 synapses per time step and 0.092 synapses per time step, respectively.

Using this model of preferential attachment, we could estimate rates of net climbing fiber synapse addition and removal that would lead to the connectivity we observed (Figures 5b and 5c). For example, if we let the synapse removal rate be 5000 times lower than the synapse addition rate, and gamma = 1.1 (these values were mid-range in the band where convergence was always reached) the simulation took 955 ± 16 time steps to reach P7-like connectivity (mean ± standard error of the mean, 100 simulations). Because the simulation covered 4 days (from P3 to P7), the average duration of a time step was thus about 6 minutes. With this information we calculated that individual climbing fiber branches added synapses at a median rate of one synapse every 7 hours and a maximum rate of one synapse every hour. In this simulation synapse removal was essentially zero over the 4 days.

### Estimating the Number of Different Olivocerebellar Axons that Innervate Developing Purkinje Cells

Above, we assessed the connectivity properties of all climbing fiber axons that innervated the fully reconstructed Purkinje cells in the P3 and P7 image volumes, where the axons were potentially split into multiple branches. Although much useful information can be revealed by studying these branches (see above), more insight still can be obtained if the number of different climbing fiber axons producing those branches is known. The direct way to get complete knowledge of the number of innervating climbing fiber axons would be to reconstruct axons back to their entry points in the cerebellar peduncle. However, this task would require collecting an image volume that is intractably large at present (millimeters on a side, >10 petabytes of raw data). Instead, we estimated the number of different climbing fiber axons innervating the fully reconstructed Purkinje cell in our P7 dataset by building on our observed climbing fiber terminal arbor morphologies and results from previous anterograde labelling. Studies using anterograde tracers showed that, in the first postnatal week, individual climbing fiber axons form multiple terminal arbors that overlap in the Purkinje cell layer (Sugihara, 2005). To calculate how many climbing fiber axons innervated one Purkinje cell, we needed to know how many terminal arbors provided innervation to that cell. However, because many of the innervating branches were incomplete, i.e. just fragments of terminal arbors, our first step was to learn the number of branch fragments a terminal arbor could supply to the volume if it were not completely within the volume. To estimate this number we spatially transposed climbing fiber terminal arbors by applying random (but constrained; see Methods) translations to the 9 complete terminal arbors in our reconstruction (Figure 6a). We then computed the number of separate branches that those transposed arbors produced by leaving and re-entering the image volume (see Methods). Using this approach we learned that a single terminal arbor is broken into 1.8 branch fragments on average in the P7 volume (Figure 6b). Thus, the 49 climbing fiber branches that innervated the fully reconstructed Purkinje cell probably came from about 27 terminal arbors. Late in the first postnatal week, an individual climbing fiber axon extends 7.4 x 10^−4^ terminal arbors/μm^2^ into each local region of cortex it innervates (computed from Sugihara, 2005; see Figures S8e-g and Methods), which, given our 9000 μm^2^ Purkinje cell layer translates to 6.7 terminal arbors per axon in the P7 volume. At P7 we estimated that ~4 different climbing fiber axons (27/6.7) innervated the fully reconstructed Purkinje cell at P7.

**Figure 6.**
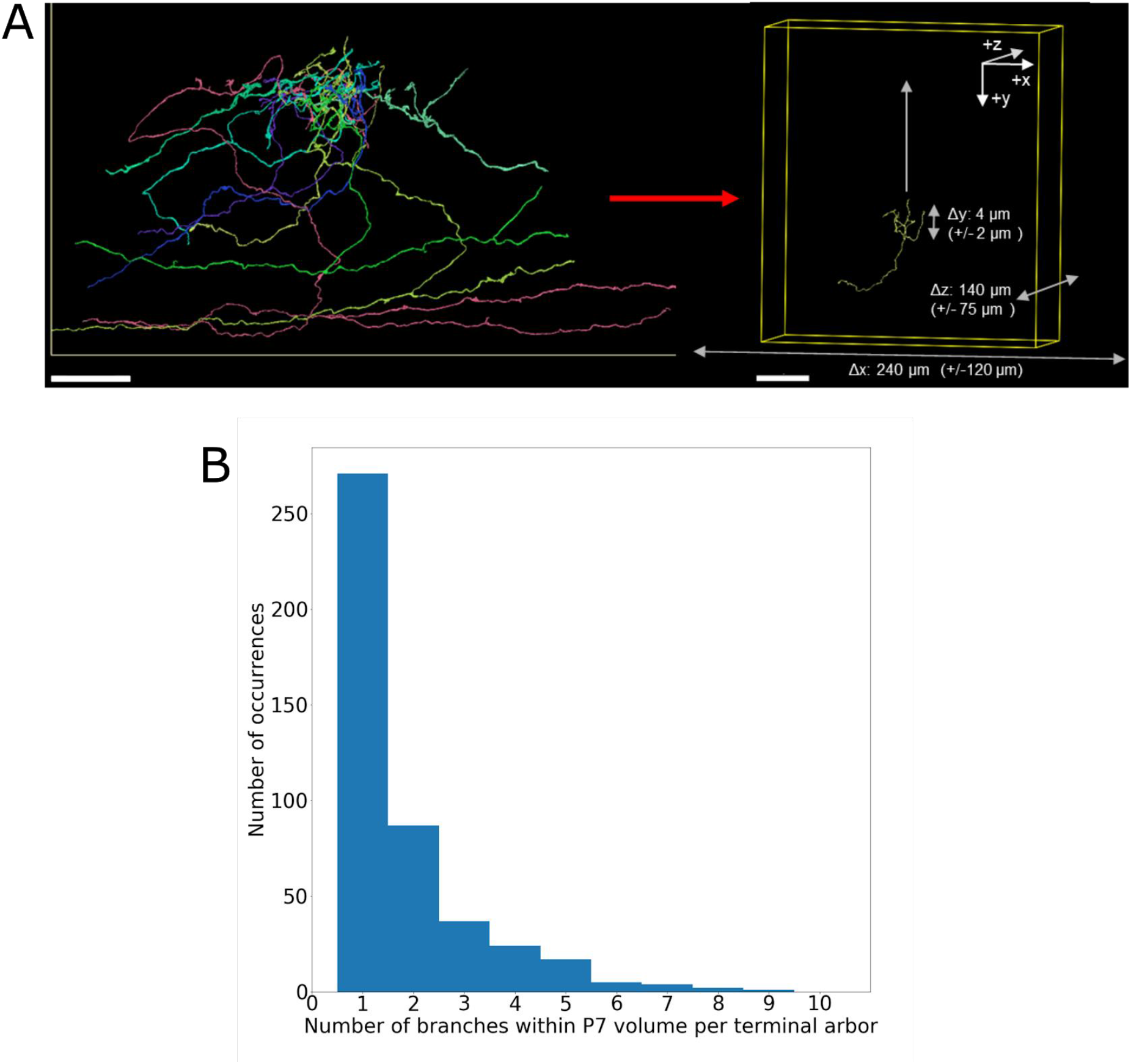
Climbing Fiber Morphology Allows Estimation of the Number of Unique Axons Innervating a Fully Reconstructed Purkinje Cell (a) Illustration showing how groups of climbing fiber terminal arbors were randomly translated, in order to estimate the number of branches that were produced by terminal arbors that left and re-entered the P7 volume multiple times. Left panel: three-dimensional rendering of the group of P7 climbing fiber branches with terminal arbors completely inside the image volume (9/49 branches). Right panel: an illustration of the bounds on translation for each selected arbor in the simulation. The upward arrow points toward the pia. The z-direction is the direction through image sections. (b) A histogram showing the number of climbing fiber branches produced by a single terminal arbor with a random displacement as described in (a). The mean of this skewed distribution is 1.8 and the standard deviation is 1.4. This plot was produced by sampling and translating terminal arbors 500 times. Scale bars: (a) 20 μm, left panel; 30 μm, right panel.

Our connectivity data also allowed us to compute how climbing fiber axons distributed their connectivity across the entire image volume. Given that 27 terminal arbors provided innervation to the fully reconstructed Purkinje cell, and given that a single terminal arbor innervated on average 4 Purkinje cells in the volume (our measurement of divergence; see Figure 3e), we estimated that 203 terminal arbors innervated the 30 Purkinje cells in the volume ((27 terminal arbors per Purkinje cell * 30 Purkinje cells)/(4 repetitions of each terminal arbor due to its divergence)). Because an individual climbing fiber forms roughly 6.7 terminal arbors in this volume (see above), we estimate that there are 30 different climbing fibers innervating the Purkinje cells in our dataset. This number corresponds to roughly one climbing fiber axon per Purkinje cell (see Discussion).

Our estimates of roughly 4 climbing fiber axons innervating a single Purkinje cell and 30 axons innervating all Purkinje cells in the volume gives us information about the divergence of individual climbing fiber axons (as opposed to any of their individual terminal arbors). For 30 axons to innervate 30 Purkinje cells such that each Purkinje cell receives 4 axon inputs, each individual climbing fiber axon should have a divergence of 4 Purkinje cells (4 axons per Purkinje cell * 30 Purkinje cells = 120 innervations of Purkinje cells by axons, from 30 axons in the volume; thus each axon innervates 120/30 = 4 Purkinje cells). Notably, this value matches the average divergence of climbing fiber branches (see Figure 3c). This information suggests that on average all branches of an individual climbing fiber axon restrict their innervation to of the same (4) Purkinje cells (or subsets thereof).

### Assessing Shared Connectivity Preferences of Branches of Climbing Fibers

After the first postnatal week, the synaptic preferences established by climbing fiber branches lead to the eventual single innervation of Purkinje cells by climbing fiber axons. Thus, a whole climbing fiber axon (not to be confused with the single branches of climbing fibers—see Figures 2c-f) must choose among its Purkinje targets which one (or ones) it will remain in contact into adulthood. The question we wished to answer is whether the branches of an individual climbing fiber are acting in concert or independently.

We used hierarchical clustering to explore the Purkinje cell preferences of the reconstructed climbing fiber branches in our P7 dataset (see Methods, Figure 7). We analyzed climbing fiber branches that showed a synaptic preference for at least one Purkinje cell (by forming at least 17 synapses at P7 on at least one Purkinje cell, i.e. greater than the 90th percentile of the synapse distributions in Figure 2d; see Figure 7a). Our clustering analysis (Figure 7b) revealed three groups of climbing fiber branches with shared preferences at P7 (https://github.com/amwilson149/baby-andross/blob/master/Neuroglancer_links/190502_p7_Group_1_2_and_3_CFs_and_syns.txt [link will be updated]). All members of each group exhibited the same preferences for a subset of Purkinje cells. In particular, the axon branches in group 1 (branches 59, 14, 7, and 15--see Figure 7c) each innervated Purkinje cell 17 more strongly than any other Purkinje cell. In total they established 105 synapses on this target cell. The same result was found for the group 2 branches (29, 4, 9, 13, 46, 5, and 37): each of those branches established the largest numbers of synapses onto Purkinje cell 25 (in total they formed 194 synapses onto that cell). The branches in group 3 (56, 3, and 72) all exhibited a synaptic preference for Purkinje cell 1 (the fully reconstructed cell; these branches established a total of 61 synapses onto that cell). The remaining climbing fiber branches that we analyzed (31, 44, 67, 2, and 33) also showed preferences, but they did not co-share preferences for single Purkinje cells as strongly as the other groups did. The shared preferences among climbing fiber branches in a group for the same Purkinje cell were statistically unlikely to have occurred by chance. If we restricted climbing fiber branches to innervate their observed Purkinje cell targets, and randomly sampled the number of synapses they formed onto each of those targets from the synapse distribution in Figure 2d (while requiring each branch to still exhibit a synaptic preference as defined above for at least one cell—see Methods), we found that branches very rarely formed groups that had preferences clustered as tightly around the same Purkinje cells as what we observed (p = 0.004; see Methods). When we repeated this analysis at P3, we did not find groups of climbing fiber branches that had significant shared Purkinje cell preferences (p = 0.36).

**Figure 7.**
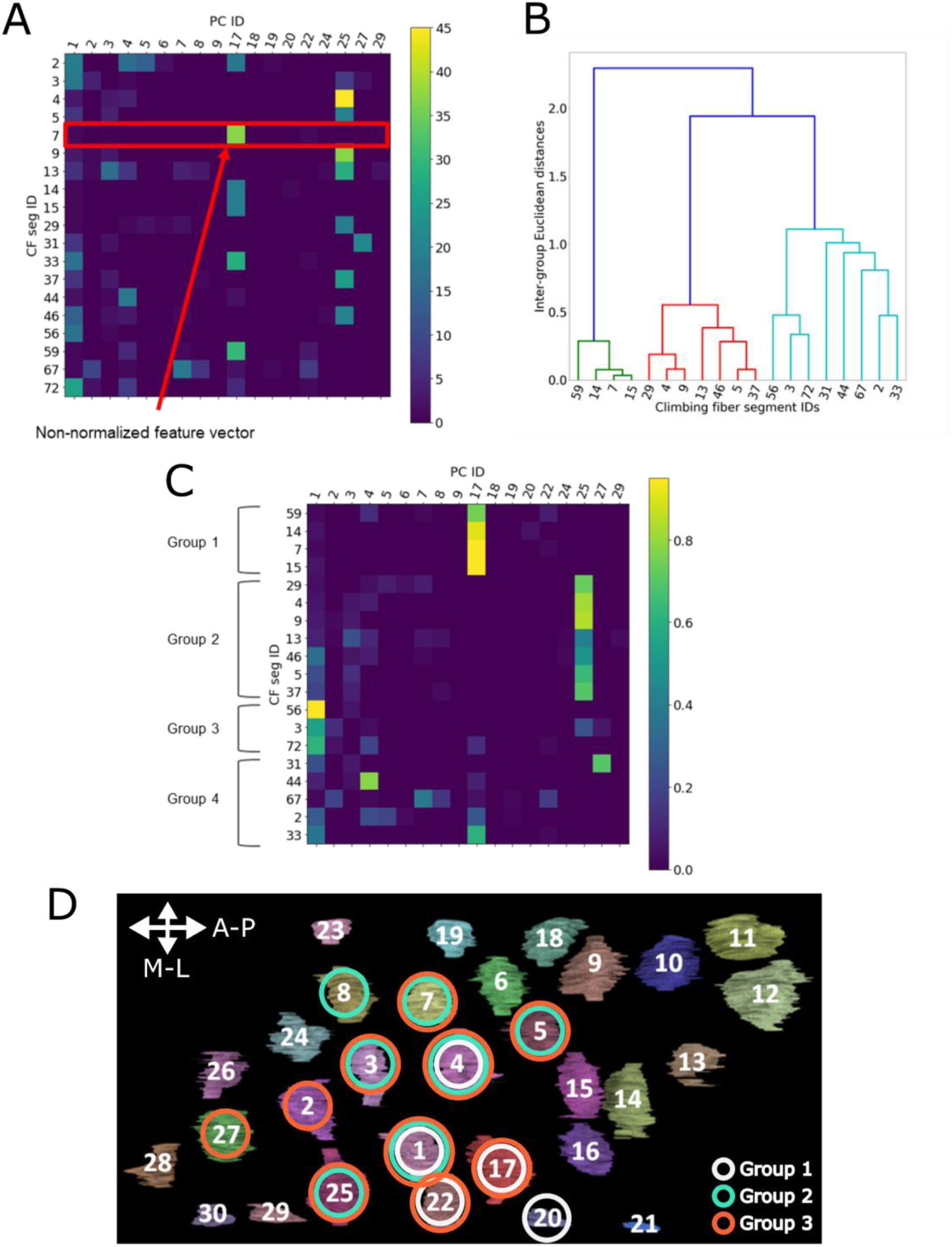
Groups of Climbing Fiber Branches at P7 have Shared Purkinje Cell Preferences (a) The P7 connectivity matrix for climbing fiber branches that exhibited strong synaptic preferences. Each row represents connectivity for a particular climbing fiber branch and each column represents connectivity for a particular Purkinje cell. The color of the element in row *i*, column *j* is the number of synapses formed by climbing fiber branch *i* onto Purkinje cell *j*. The connectivity for a given climbing fiber branch (red box) can be interpreted as a feature vector in Purkinje cell space, where the number of synapses formed onto each Purkinje cell is the length of the vector component in that dimension. (b) Dendrogram showing the groupings of the P7 climbing fiber branches in a hierarchical clustering process. Branches with the most similar Purkinje cell connectivity (i.e. most similar feature vectors) are grouped together at earlier stages (i.e. lower levels of the dendrogram). The height of the bars that link two groups of climbing fiber branches give the distance between the feature vectors of those two groups (i.e. shorter bars indicate more similar groups). (c) The observed P7 connectivity matrix, reordered to match the similarity-based ordering prescribed by hierarchical clustering in (b) (so that branches with the most similar connectivity are nearest to one another). Groups 1, 2, and 3 contain climbing fiber branches that share preferences for the same Purkinje cells. Group 4 contains branches that did not share preferences for Purkinje cells with other branches included in the analysis. (d) The somas of Purkinje cells in the P7 volume (viewed along the radial direction, looking from the pia to the white matter; arrows indicate the anteroposterior and mediolateral axes). Preferred Purkinje cells for each group of branches are identified using circles (gray for group 1, cyan for group 2, and orange for group 3). The number on each Purkinje cell is its segment ID.

Thus, as climbing fiber branches form synapses over the first postnatal week, groups of them develop preferences for the same sets of Purkinje cells. These preferences do not appear to be a result of spatial factors, as we could find no topographical patterns to the sets of preferred cells for each climbing fiber group we identified at P7 (Figure 7d).

The climbing fiber branches with shared Purkinje cell preferences at P7 may be related in two ways: they might be branches of the same climbing fiber axon, or they might be branches from different climbing fiber axons. We considered the likelihood of these relationships by again considering data about climbing fiber morphology and divergence. First, we calculated that the 49 climbing fiber branches innervating the fully reconstructed Purkinje cell corresponded to approximately 4 climbing fibers. Thus, we expect that the 19 branches for which we have assessed Purkinje cell preferences originate from a smaller number of different climbing fibers (i.e., ~4 or fewer because all these branches innervated the fully reconstructed Purkinje cell).

For branches of the same axon, we hypothesize that their synaptic preferences are similar, i.e., they fall into the same groupings (see above). If they did not fall into the same groups, then two things must occur to generate the results in Figure 7c: first, different branches of the same axon would have to exhibit different synaptic preferences; and second, subsets of branches from different axons would have to share the same synaptic preferences. We cannot conceive of a means of dividing branches in this way. Moreover, electrophysiological studies consistently show that, at this developmental age, only one climbing fiber axon (among the multiple inputs) is strong (Kano et al., 2018) which contradicts the idea of multiple axons having the same synaptic preferences.

These results lead to the conclusion that many branches of the same axon have similar synaptic preferences. We believe that, because of this feature for branches of the same axon, it is possible to infer the origin axon for different branches based on their synaptic connectivity alone. This conclusion has important implications not only for the development of the cerebellum but for connectomics in general (see Discussion).

## DISCUSSION

In this study, we produced electron microscopy image volumes of mouse cerebellum at P3 and P7 and reconstructed climbing fiber branches and their Purkinje cell targets, in order to learn more about the changing organization of climbing fiber input to Purkinje cells during the first postnatal week. Our results reveal several features of climbing fiber-Purkinje cell synapse elimination that were not previously known. First, during this time, single climbing fibers form many additional synapses to focus their innervation onto a subset of their Purkinje cell targets. Second, we find that single-synapse strengths (as measured by PSD area) become more uniform and do not correlate with the overall strength of a climbing fiber-Purkinje cell connection. These two points, combined with our confirmation that synaptic pruning does not occur, lead us to conclude that addition of synapses is the primary factor underlying climbing fiber functional differentiation in the first postnatal week. This structural information is more consistent with the electrophysiological evidence of synaptic strengthening during the first postnatal week (Hashimoto and Kano, 2003; Bosman et al., 2008) as opposed to results suggesting synaptic changes begin in the second postnatal week (Scelfo and Strata, 2005). Third, we infer that the synapse addition between P3 and P7 involves positive feedback between climbing fibers and Purkinje cells. Using simulations we found that a “rich get richer” strategy of climbing fibers adding progressively more synapses onto Purkinje cells they already strongly innervated was the only way P3 connectivity could evolve into P7 connectivity (Figure 5). Furthermore, at both P3 and P7 the synapse distributions (Figure 2b) were well fit by power laws, and distributions following a power law are a known result of positive-feedback-based addition, or preferential attachment (Newman 2005).

We also addressed a challenge that beset our analysis and is present in many other connectomic image volumes, namely, that practically all axons are incomplete because connectomic volumes are small relative to neuron sizes. We combined information about large-scale climbing fiber morphology (Sugihara, 2005) with our own reconstructions of terminal arbors to quantify the total number of climbing fiber axons that provided innervation to the 30 Purkinje cells in our P7 image volume. We believe this is the first time that the number of axons innervating a local region of brain has been quantified. This information provides important context about climbing fiber-Purkinje cell synapse rearrangement. We found, interestingly, parity, in that there are 30 climbing fibers innervating 30 Purkinje cells. Because each Purkinje cell is innervated by 4 climbing fibers on average (see Figure 6), the maximum number of climbing fibers that could have provided innervation to this volume was ~120 (i.e. 4 distinct axons per Purkinje cell x 30 Purkinje cells). The minimum number of climbing fibers that could have provided 4 inputs to this set of Purkinje cells is 4 (if the same 4 axons innervated every Purkinje cell). The actual number of inputs, 30, is exactly what one would expect to find innervating a volume of 30 Purkinje cells in adult cerebellum, because adult climbing fibers innervate Purkinje cells sparsely; they rarely innervate more than one cell in a field of 30. More specifically, an adult climbing fiber forms synapses with ~10 Purkinje cells out of thousands in a few lobules of the cerebellum (Sugihara, 2001; Fujita and Sugihara, 2013). Thus, climbing fiber branching in development appears to be economical: the number of axons that innervate Purkinje cells within a local region in the first postnatal week is just enough to assure that each axon ends up with a postsynaptic target and that none branched there in vain (Figure 8).

**Figure 8.**
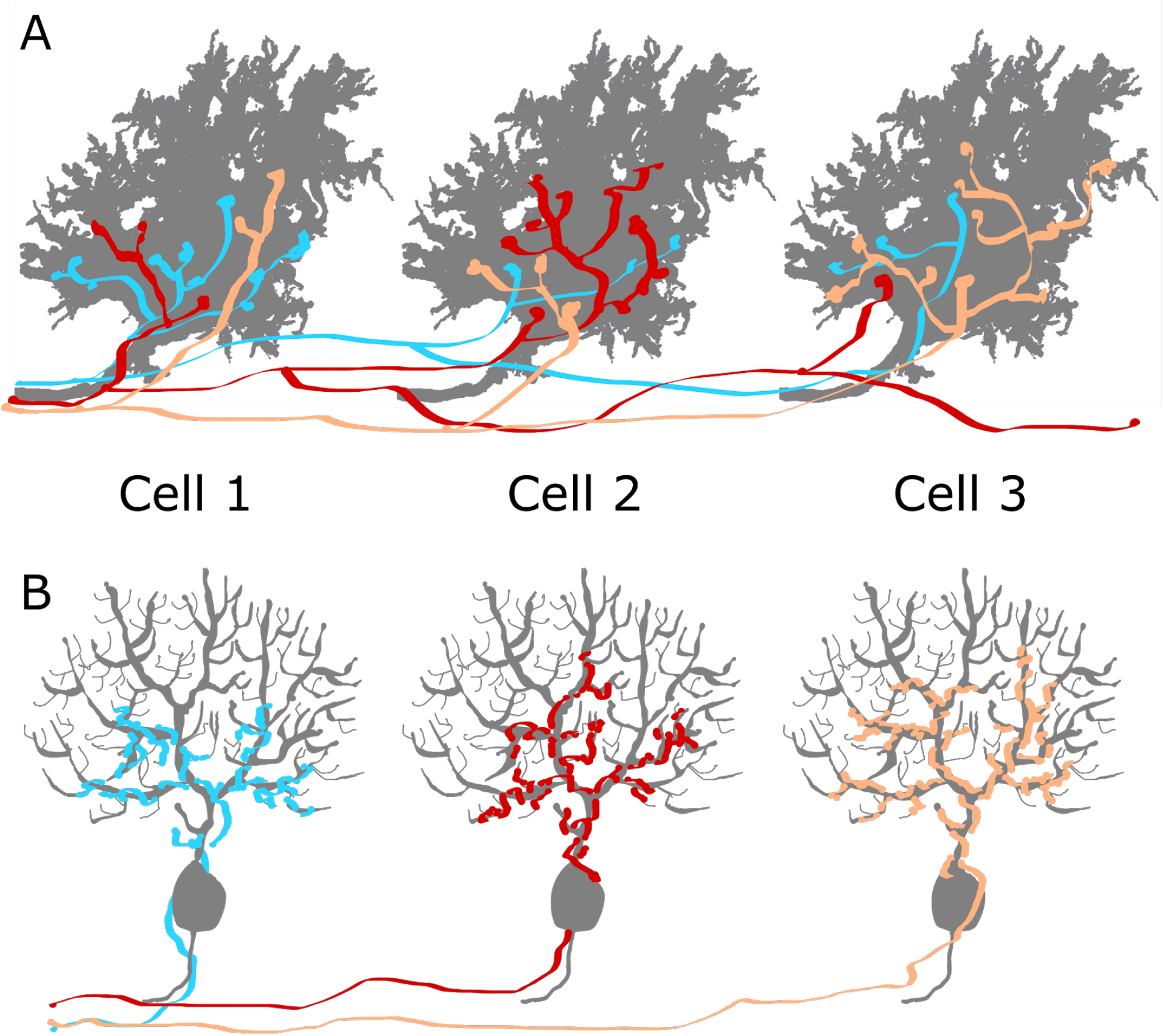
Climbing Fiber-Purkinje Cell Synaptic Rearrangement during Postnatal Development is an Economical Process (a) Illustration demonstrating the observed pattern in Purkinje cell innervation by climbing fiber axons late in the first postnatal week. In a local region, the number of climbing fiber axons roughly matches the number of Purkinje cells in the region. Each climbing fiber axon develops a synaptic preference for a small number of Purkinje cells (e.g. Cell 2 for the red climbing fiber axon illustrated here) by adding synapses to those cells, while remaining synaptically connected to non-preferred cells (for the red axon, Cells 1 and 3). (b) Illustration showing the schematic described in (a) after maturity is reached. Each climbing fiber has remained synaptically connected to one Purkinje cell in the local region (e.g. the red axon has remained connected to Cell 2) and has removed its synapses from non-preferred Purkinje cells (Cells 1 and 3 for the red axon). Each axon forms the hundreds of synapses observed between single climbing fiber-Purkinje cell pairs in adulthood, and ramifies along the proximal dendrites of its target Purkinje cell. The number of climbing fiber axons in a local region of cerebellar cortex still roughly matches the number of Purkinje cells.

This information, taken together, provides a glimpse at how development translates into structural changes in brain circuitry. It will also be useful for constraining future mechanistic models of cerebellar development: any model should produce circuitry changes consistent with those revealed in this analysis.

Finally, although we studied synaptic connectivity from climbing fiber branches onto Purkinje cells, these electron microscopy volumes also contain ample information about all other types of cells found in the cerebellar cortex, along with their intracellular organelles and their synaptic connectivity. These image volumes are thus useful resources for investigations of normal cerebellar development in mouse. The volumes and our annotations of them are available at https://bossdb.org/project/PREwilson2019 [link will be updated].

### Branches of the Same Climbing Fiber Exhibit Similar Synaptic Preferences via a Contact-Mediated Mechanism

Our analysis indicates that different climbing fiber branches in the P7 volume had statistically significant similarities in their synaptic connectivity. These similarities were not explained by close fasciculation of these axon branches. Indeed, each climbing fiber branch appeared to have a completely individual branching pattern (https://github.com/amwilson149/baby-andross/blob/master/Neuroglancer_links/190502_p7_Group_1_2_and_3_CFs_and_syns.txt [link will be updated]). Despite these differences in shape, a number of these branches could be grouped by remarkably similar shared synaptic preferences for Purkinje cells (Figure 7c). Based on a number of arguments (see Results) we think it likely that each such group of climbing fiber branches originates from one olivocerebellar neuron’s axon. This finding leads to an important implication for the way that preferred Purkinje cells are chosen. Namely, these axons do not establish synaptic preferences with a particular Purkinje cell through specific, directed arborization during the first postnatal week. Rather, they establish large numbers of synapses at sites where they happen to be in contact. In this sense the preferences are contact-mediated rather than axon growth-mediated. This idea is strengthened by the observation that climbing fiber branching is not pronounced in the vicinity of preferred cells compared with other Purkinje cells. Rather, the only evidence of the site of the preferred Purkinje cell is the greater density of synapses per axon length (see https://github.com/amwilson149/baby-andross/blob/master/Neuroglancer_links/190502_p3_CFs_and_syns_MFs_and_syns_PC1_and_all_PC_somas.txt at P3, and https://github.com/amwilson149/baby-andross/blob/master/Neuroglancer_links/190502_p7_CFs_and_syns_MFs_and_syns_PC1_and_all_PC_somas.txt at P7 [links will be updated]). One interpretation of this situation is that between P3 and P7 axons add synapses along existing branches that are already juxtaposed with the preferred Purkinje cell target.

This finding also illustrates an important point for connectomics datasets in general: namely, it may be possible to regroup broken axon pieces by leveraging their synaptic connectivity, as we have done for the P7 cerebellum dataset. This strategy should allow for more complete axonal reconstructions and therefore more accurate connectivity analysis. This type of connectivity-based inference is only possible with analysis in high-resolution connectomics datasets, in which all synapses formed by branches of an axon can be identified and their distributions across targets can be directly measured.

### Functional Differentiation in the Cerebellum vs. in the Neuromuscular Junction

Our observations indicate that, in the first postnatal week, climbing fiber axons develop preferences for certain Purkinje cells as they add synapses (Figures 7b and 7c; see previous section). Based on the idea that multiple branches originate from the same axon (see Figure 6 and Results) there may be hundreds of synapses added by one axon onto one Purkinje cell between P3 and P7 while other axons innervating the same Purkinje cell change their synaptic input much more modestly. Given the loss of all but one input over next few weeks, it seems likely that the axon with the large increment in synapses will be the remaining climbing fiber after synaptic reorganization is complete. Importantly, there was no evidence in our studies of synapse loss during the first postnatal week. In the second and third postnatal weeks, weaker climbing fiber inputs do lose all their synapses with a Purkinje cell. This connection loss leaves Purkinje cells singly innervated by one dominant climbing fiber that adds additional synapses as it climbs up the Purkinje cell’s dendritic tree.

This sequence of events for climbing fibers in the cerebellum contrasts with developmental synapse rearrangement of motor axons at the neuromuscular junction, although in both cases postsynaptic targets are initially innervated by multiple axons and end up with a single axonal input. At the neuromuscular junction, the multiple axons that converge undergo a process of synapse exchange: an axon adds synaptic territory by taking over space that was vacated by a different axon (Walsh and Lichtman, 2003; Gan and Lichtman, 1988; Turney and Lichtman, 2012). Synapse addition and removal are therefore inextricably linked. The significance of this difference is that at the neuromuscular junction and certain other sites (Chen and Regehr, 2000; Lichtman, 1980) synapse loss appeared to be a requirement for the strengthening of the ultimate surviving input. In the cerebellum, electrophysiological evidence shows relative strengthening of one climbing fiber input compared with others in the first three postnatal weeks (Kano et al., 2018) and is consistent with the profound synaptogenesis we describe here. However, until now it had not been possible to know whether climbing fiber strengthening in the cerebellum also requires concurrent synapse removal by other climbing fiber inputs. Our results provide evidence that synaptic strengthening of climbing fiber inputs is unrelated to synapse removal by other climbing fibers. In particular axonal branches are not pruned (Figure 3) and the evolution of the P3 to P7 innervation patterns requires that no synapses are removed (Figure 5). A classical Hebbian mechanism (i.e. strengthening without elimination) may underlie the establishment of a dominant input in the cerebellum (Hebb, 1949; see also Lichtman and Balice-Gordon, 1990).

### Persistence of Weak Climbing Fiber Inputs

The persistence of weak climbing fiber inputs despite the emergence of a dominant input during the first postnatal week raises the question of why these neurons would maintain weak connectivity. In particular, whereas a climbing fiber axon seems to form hundreds of synapses onto a preferred Purkinje cell, it can form very few synapses onto cells where it is a weak input. For example, the branches in group 1 of our preference analysis (Figure 7c), which likely stem from a single axon, collectively form 105 synapses onto Purkinje cell 17, their most preferred target, but form only 4 synapses onto Purkinje cell 1, which is a non-preferred target. Physiologically effective climbing fiber axons in an adult establish many hundreds of synapses with each of their target cells, so it is unlikely that axons forming only a few synapses have functional significance. One possible reason for maintaining weak connections is that it provides axons with a foothold on a target cell should the dominant input be damaged during development (see for example Carrillo et al., 2013; Turney and Lichtman, 2012), or in case interactions between climbing fibers and Purkinje cells require a dominant input to refocus its resources elsewhere as has been proposed at the developing neuromuscular junction (Walsh and Lichtman, 2003). In this sense, Purkinje cells may be hedging their bets, so to speak, by remaining connected to multiple climbing fibers before the competition is completely resolved. From the perspective of the climbing fiber the same may be true: an axon may remain connected to many target cells to assure that it still innervates a few after most synaptic pruning has occurred.

### Comparative Connectomics

Connectomics per se is a descriptive approach. It relies on inductive reasoning, so that hypotheses are generated more easily than tested. One way, however, to generate and test hypotheses in connectomics data sets is to compare samples that differ in some way. In this study we have compared connectomics data from two developmental stages to learn how neural circuits become modified in early postnatal life. The power of this strategy is that a fundamentally static technique (looking at stained postmortem tissue) can be used to infer information about a highly dynamic phenomenon (the maturation of neural circuits). One challenge is that comparing connectomes is still a nascent approach. Although we possess potentially vast amounts of structural data from two time points, we do not yet know what are the ideal ways to make statistically rigorous comparisons. We contend that these two data sets contain material for an essentially unlimited number of hypotheses about how the cerebellum changes over the first postnatal week. Determining the best way to extract the things that are different and the things that aren’t from multiple samples will require turning these “digital tissues” into systematized databases that can be scrutinized automatically. This will likely be one of the central thrusts of connectomics going forward.

## Supporting information

All Supplemental Figures and Tables

## AUTHOR CONTRIBUTIONS

Conceptualization, A.M.W. and J.W.L.; Methodology, A.M.W. and J.W.L.; Software, A.M.W, A.S.-P., T.R.J., S.K.-B., and H.P.; Formal Analysis, A.M.W. and J.W.L.; Investigation, A.M.W.; Resources, J.W.L. and R.S.; Data Curation, A.M.W., A.S.-P., T.R.J., S.K.-B., and H.P.; Writing - Original Draft, A.M.W. and J.W.L.; Writing - Review & Editing, A.M.W. and J.W.L.; Visualization, A.M.W. and J.W.L.; Supervision, J.W.L.; Funding Acquisition, A.M.W. and J.W.L.

## ACKNOWLEDGMENTS

We thank Daniel Berger, Daphna Keidar, and Will Silversmith for helpful discussions, and thank those who helped with annotating the datasets (Silvia Caminiti, Lena Jiang, Jasmine Lopez, Brenda Marin-Rodriguez, Molly McGowan, Renuka Nannapaneni, Bryan Nelson, Stephen Ng, and Vinutna Veeragandham). We gratefully acknowledge support from the NIH/NINDs (National Research Service Award F31NS089223,1DP2OD006514-01, TR01 1R01NS076467-01, and 1U01NS090449-01), the National Defense Science and Engineering Graduate Fellowship (NDSEG) Program, Conte (1P50MH094271-01), MURI Army Research Office (contract no. W911NF1210594 and IIS-1447786), NSF (OIA-1125087 and IIS-1110955), the Human Frontier Science Program (RGP0051/2014), the NIH and NIGMS via the National Center for Multiscale Modeling of Biological Systems (P41GM10371), and IARPA contract D16PC00002.

## METHODS

### Data Acquisition

All animals were handled according to protocols approved by the Institutional Animal Care and Use Committee at Harvard University.

We prepared samples from the cerebella of two CD1 wild-type unsexed mouse pups, one aged P3 and one aged P7, from timed-pregnancy mothers. Cages were checked twice a day for pups (once in the morning and once in the evening) and P0 was assigned at the time when pups were found. Ages are thus accurate to within 12 hours. We anesthetized pups with 0.01 mL of sodium pentobarbital (50 mg/mL) and intracardially perfused each with 2%-2% paraformaldehyde/glutaraldehyde in 0.15-M sodium cacodylate, 2-mM CaCl_2_ buffer solution. We qualitatively monitored pup behavior to ensure that they behaved similarly to their littermates, and we measured brain sizes along the anteroposterior, mediolateral, and dorsoventral axes to ensure they fell within normal ranges. We then isolated the cerebellum from each sample and cut it into 300-μm-thick parasagittal sections. From each cerebellum, we chose one thick section in the medial part of the vermis and stained it using a ROTO protocol (Willingham and Rutherford, 1984) (2% osmium tetroxide plus 0.015 g/mL potassium ferrocyanide, 1% thiocarbohydrazide, and 2% osmium tetroxide). We then dehydrated and embedded each section in Epon 812 resin. We cut the embedded tissue into 30-nm sections using an automatic tape collecting ultramicrotome (ATUM) and post-stained each ultrathin section with 4% uranyl acetate and 4% lead citrate (Kasthuri et al., 2015).

We imaged these sections using secondary electron detection in a single-beam scanning electron microscope (1.7 kV; ZEISS Sigma). The acquisition was automated using WaferMapper software (Hayworth et al., 2014). Images were acquired at a resolution of 4 nm x 4 nm per pixel at a dwell time of 200 ns per pixel.

We reconstructed a Purkinje cell roughly in the middle of the P3 volume and one roughly in the middle of the P7 volume. In each case we also reconstructed all of its synaptic inputs. For each of the innervating axons we identified all of their other synapses and postsynaptic partners in the volume. We also reconstructed the somata of all the Purkinje cells in the volume at P3 and P7. Most of this reconstruction was done by computer assisted manual tracing using the VAST-lite tool (Berger, Seung and Lichtman, 2018). In the P7 volume we began with a machine learning based segmented subvolume (72 μm x 72 μm x 30 μm) generated by Rhoana (Kaynig et al., 2015) and edited it using Mojo software (Seymour Knowles-Barley, http://vcg.seas.harvard.edu/presentations/mojo-20-connectome-annotation-tool). The axon branches and cells in the subvolume were then extended throughout the volume using VAST.

Because our goal was to analyze the climbing fiber input to Purkinje cells it was important to classify each presynaptic axon branch in order to exclude axons that were not climbing fibers. We therefore determined the ultrastructural characteristics of the synapses it formed (excitatory vs. inhibitory) to rule out inhibitory axons (i.e., Purkinje cell collaterals, stellate, basket, Golgi, Lugaro, and candelabrum cells; Palay and Chan-Palay, 1974; Lainé and Axelrad, 1994). Inhibitory axons were identified by their lack of a pronounced postsynaptic density, a feature of excitatory synapses (Peters, Palay, and Webster, 1991). Inhibitory synapses also possessed irregularly shaped vesicles (unlike the round vesicles we observed in excitatory axons). If a synapse was ambiguous we traced the axon to one or more additional synapses and this allowed an unambiguous determination of whether it was excitatory or inhibitory. We also could rule out excitatory granule cell axons by their distinctive morphology (i.e., a smooth, unbranched axon segment that ran parallel to many other similar axon segments along the mediolateral axis in the molecular layer, a feature of the parallel fiber portion of a granule cell axon; see Palay and Chan-Palay, 1974). We also excluded any axon that could be traced back to its cell body because climbing fibers originate outside the cerebellum, in the inferior nuclear olive (this allowed us to classify and remove inputs that were the radial ascending branch of a granule cell). The remaining excitatory axon branches were either climbing fibers, or mossy fibers (which originate in various nuclei outside the cerebellum as well as from recurrent collaterals of excitatory cells in the deep cerebellar nuclei inside the cerebellum, and from unipolar brush cell axons within the cerebellar cortex; see Hess, 1982; Mugnaini and Floris, 1994; and Mugnaini, Sekerkova, and Martina, 2011).

Because mossy fibers have been reported to make transient connections on Purkinje cells in development (Mason and Gregory, 1984; Kalinovsky et al., 2011), it was important to exclude these excitatory inputs to Purkinje cells as well. We therefore analyzed a subset of excitatory (non-granule cell) axon branches to identify all of their postsynaptic targets at both P3 (20 branches) and P7 (14 branches). We did this painstaking analysis in order to learn whether mossy and climbing fibers had different connectivity profiles. At each age we computed the fraction of synapses formed by these axon branches onto (1) Purkinje cells, (2) granule cells, and (3) interneurons. At both ages we found a clear splitting of branches into two groups (Figure S2): branches that formed a much larger fraction of their synapses onto Purkinje cells relative to granule cells, and branches that formed a much larger fraction of their synapses onto granule cells relative to Purkinje cells.

Based on this preliminary result, we attempted to find a quantitative criterion to split the climbing fibers from the putative mossy fibers. We therefore wrote code to perform two modes of cluster analysis (k-means and hierarchical agglomerative clustering--https://github.com/amwilson149/baby-andross/blob/master/181114_partition_P3_P7_inputs_w_k-means_DBSCAN_Agglom.ipynb) to group the branches we analyzed based on their connectivity properties. We also included a set of axons that we believe are definitely mossy fibers (12 at P3 and 10 at P7) because they innervate the dendrites of granule cells deep in the internal granular layer. Both clustering methods produced nearly the same classification metric (see Figure S2). Based on this analysis, the axon branches could be completely separated. At P3 axon branches that formed at least 40% of their synapses with Purkinje cells (they formed the rest with interneurons and rarely with granule cells) were a distinct class from the other axons that we sampled which made larger numbers of synapses with granule cells and very few synapses with Purkinje cells. This latter group clustered with the actual mossy fibers. Thus we think that this clustering allowed us to exclude mossy fibers from our climbing fiber population. At P7 the same approach clustered the climbing fibers by choosing branches that formed at least 70% of their synapses on Purkinje cells. We excluded any axon that established fewer than 5 synapses in the volume from our analysis.

The P3 and P7 image volumes and segmentations are freely available at https://bossdb.org/project/PREwilson2019 [link will be updated].

### Power Law Fits

Fits were computed for the synapse distributions (number of synapses formed by single climbing fiber branches onto each Purkinje cell they innervated) using the scipy.optimize.curve_fit function in Python.

### Skeletonization for Climbing Fiber Branch Length and Synapse Density Measurements

Skeletons were computed for climbing fiber branches in both datasets using Kimimaro (https://github.com/seung-lab/kimimaro) and Igneous (https://github.com/seung-lab/igneous) software and can be viewed in Neuroglancer (https://github.com/amwilson149/baby-andross/tree/master/Neuroglancer_links [link will be updated]). The total length of each branch was queried with this software using the cable_length attribute for these skeletons. To compute the length of each climbing fiber branch that was in the Purkinje cell layer, we cropped the skeletons to include only the pieces above the Purkinje cell layer-internal granular layer interface (i.e. the pieces in the upper 75% of the volume at each age) and then computed the cable length. Synapse densities in the Purkinje cell layer were computed by dividing the total number of synapses formed by a climbing fiber branch onto Purkinje cells in the Purkinje cell layer by the total length of the branch in the Purkinje cell layer.

### Estimation of Postsynaptic Density Area

We manually traced the PSDs of all the climbing fiber segment-Purkinje cell synapses using VAST. To accurately represent PSD shapes with these annotations, we converted voxel lists from pixel values (which were anisotropic: 4 nm/px x 4 nm/px x 30 nm/px) to units of nm. We then performed principal component analysis on each voxel list and used the third component to define the normal to the flattest plane of the PSD. We then projected the PSD voxel list onto this plane to create a 2-dimensional version of the PSD. We computed the convex hull of the projected pixels and used it to compute an estimate of the PSD area. This method was insensitive to factors like the orientation of a PSD within the image volume.

### Time-Evolution Simulation

We wrote a simulation to evolve synaptic connectivity matrices using stochastic synapse addition and removal in order to explore the types of processes that could generate the connectivity observed at P7, given an initial connectivity identical to what we observed at P3. In the simulation the connectivity matrix was initialized to the observed P3 connectivity (i.e. 55 rows x 30 columns with values producing the distributions in Figure 2c). A target connectivity matrix was set to the observed connectivity at P7 (49 rows x 18 columns with values producing the synapse distribution in Figure 2d). For both ages, the first column represented the connectivity by climbing fiber branches onto the fully reconstructed Purkinje cell.

In each timestep, we (1) removed a synapse from several randomly chosen connected climbing fiber branch-Purkinje cell pairs (i.e. we decremented the values of several nonzero connectivity matrix elements). We then (2) compressed the connectivity matrix by (a) removing climbing fiber branches that became completely disconnected from the Purkinje cells in our dataset (rows containing only zeros), (b) removing Purkinje cells that became completely disconnected from the climbing fiber branches in our dataset (columns containing only zeros), and (c) removing climbing fiber branches that were no longer synaptically connected to the fully reconstructed Purkinje cell (rows for which the element in the first column became zero). We removed the rows that met condition (c) so that the connectivity matrix always reflected what we would have measured by reconstructing circuitry as we did in our P3 and P7 datasets (i.e. by identifying the climbing fiber synaptic inputs to one fully reconstructed Purkinje cell and then adding connectivity information for all the other Purkinje cells they innervated in the volume). We then (3) added synapses at several randomly chosen climbing fiber segment-Purkinje cell interfaces (i.e. incremented the value of some nonzero matrix elements). Finally, we (4) compared the resulting connectivity matrix with the target matrix to determine whether the two were statistically indistinguishable (see below). If they were, we regarded this time step as one in which P7-like synaptic connectivity had been reached. The software we developed for this purpose can be found here (along with a description of the parameters): https://github.com/amwilson149/baby-andross/blob/master/190131_evolve_synaptic_conn_p3_p7.py.

To determine whether a simulated connectivity matrix was statistically indistinguishable from the target matrix, we compared the two using four features: (1) the full synapse distribution (number of synapses per climbing fiber branch onto each of its Purkinje cell targets), (2) the distribution of total number of synapses formed by each climbing fiber branch (row-wise weight distribution), (3) the distribution of the number of Purkinje cells innervated by each climbing fiber (i.e. climbing fiber divergence, or row-wise weight density), and (4) the distribution of the number of climbing fibers innervating each Purkinje cell (i.e. climbing fiber convergence onto Purkinje cells, a measure of column-wise weight density). We also tracked the sizes of the matrices (i.e. the number of climbing fiber branches (rows) and the number of Purkinje cells (columns)) and required them to not become more different than the size difference between P3 and P7. However, the sizes could be derived from (1) and (2) above, so these measurements were not independent constraints on the simulation. Rather, we used them as a sanity check.

We considered properties (1) through (4) for the simulated and P7 matrices to be statistically indistinguishable if a Wilcoxon rank sum test failed to detect a difference at an alpha level of 0.05. We required all of these properties to be statistically indistinguishable for the simulated matrix to be considered as having reached the actual P7 configuration.

### Estimating the Number of Branches Produced By a Climbing Fiber Terminal Arbor

To estimate the number of branches that a climbing fiber terminal arbor extended into our image volumes, we used our reconstructions of unbroken climbing full terminal arbors at P7 (9/49 branches) as templates and simulated how similar arbors could be broken by crossing the image volume boundaries. For each simulation we randomly selected a terminal arbor from this reconstructed set with replacement. We then applied randomly chosen but constrained translations to the voxel list for that terminal arbor. Within the Purkinje cell layer (i.e. the x-z plane), we allowed translations in any direction by any amount (with uniform probability) that ranged from zero up to a displacement of 120 μm (the volume width) in x and up to 75 μm (the volume depth) in z. A terminal arbor at the maximum displacement in either dimension would thus be located just outside the volume. We also allowed translations in either direction along the y-axis of up to 2 μm to allow varying arbor positions while enforcing the biological constraint that climbing fiber terminal arbors should terminate in the Purkinje cell layer. We then applied the bounding box of the image volume ((0 μm, 0 μm, 0 μm) to (120 μm, 190 μm, 75 μm)) to the resulting voxel list. Specifically, any voxel of a simulated terminal arbor that fell outside the image volume was removed, to reflect the fact that that portion of the arbor would not have been reconstructed. For the remaining voxels we then counted the number of connected components. We repeated this process 500 times to produce a distribution of the number of branches produced by broken terminal arbors in the P7 image volume.

### Estimating the Density of Terminal Arbors per Climbing Fiber Axon in the First Postnatal Week

We computed the number of terminal arbors extended per climbing fiber axon in a single microzone of cerebellar cortex (usually a part of a lobule) during the first postnatal week using measurements reported in Figures 1b and 6a of (Sugihara, 2005). This data contained images of dye-labeled climbing fiber axons and the majority of their terminal arbors, according to the report. We estimated densities of 8.7 x 10^−4^ terminal arbors/μm^2^, 9.8 x 10^−4^ terminal arbors/μm^2^, and 2.99 x 10^−4^ terminal arbors/μm^2^ in lobules II, III, and crus Ic, respectively, from Fig. 1B and 8 x 10^−4^ terminal arbors/μm^2^ in lobule VIb-c from Fig. 6A, in the transverse plane of the cerebellar cortex (i.e. the plane parallel to the Purkinje cell layer). The area in the transverse plane of the P7 volume is 9000 μm^2^. A single olivocerebellar axon with average terminal arbor density (7.4 x 10^−4^ terminal arbors/μm^2^, computed from the estimates above) that fully overlapped the P7 image volume would thus extend 6.7 terminal arbors into the volume.

### Matrix Reordering to Group Climbing Fiber Axon Branches

We attempted to group climbing fiber branches by the similarity of their synaptic connectivity onto the Purkinje cells in the volume at P3 and P7. We restricted our analysis to those branches that had strong preferences for at least one Purkinje cell. We considered a climbing fiber branch to have a strong preference if it formed a number of synapses onto at least one cell that was in the top 10th percentile of the connection sizes we observed (i.e. more than 3 synapses at P3 and more than 16 synapses at P7; Figures 2c and 2d, respectively). This population consisted of 23 climbing fiber branches at P3 and 19 branches at P7. For this population of climbing fiber branches, we treated the connectivity for each branch onto the Purkinje cells they innervated (a row of the connectivity matrix) as a feature vector, and computed the Euclidean distance matrix for the normalized feature vectors. The (*i*,*j*)th element of the distance matrix represents how different the connectivity onto all Purkinje cells is for climbing fiber branches *i* and *j*. We then used hierarchical agglomerative clustering (see, e.g. James et al., 2013, Chapter 10) to group the climbing fiber branches according to this distance metric.

### Significance Testing of Climbing Fiber-Purkinje Cell Connectivity Patterns

We used Monte Carlo simulations, resampling tests, and permutation tests to determine whether the various properties we observed could have resulted from random synaptic connectivity. These properties were (1) the distributions of the number of synapses formed by climbing fiber branches onto individual Purkinje cells in each dataset, (2) the correlations between PSD area of a synapse and the total number of synapses formed by its presynaptic climbing fiber branch onto its postsynaptic Purkinje cell, and (3) the ordering of climbing fiber branches into groups with shared Purkinje cell preferences, based on the numbers of synapses they formed onto each Purkinje cell target in our dataset.

For (1) the synapse number distributions, we performed a Monte Carlo simulation of the case where each climbing fiber branch formed synapses onto all its Purkinje cell targets with uniform probability. In each iteration of the simulation, we held the total number of synapses formed by each climbing fiber branch fixed at observed values, and required that the climbing fiber branch could only form synapses onto its observed Purkinje cell targets. For each synapse formed by the climbing fiber branch we chose the identity of the Purkinje target from the set of observed Purkinje cell targets, with replacement, according to a uniform probability distribution. We then computed the median and skewness of the resulting connectivity distribution. We repeated this simulation 100,000 times and compared the resulting median and skewness distributions with the observed values by computing p-values at a significance level of 0.05 (Figure S5).

For (2) the PSD area correlations, we performed a permutation test to simulate how PSD area would relate to number of synapses for single climbing fiber branch-Purkinje cell pairs if they actually had a random relationship. In each iteration, we randomly permuted PSD areas for all climbing fiber branch-Purkinje cell synapses and computed the Pearson’s correlation coefficient. We repeated this procedure 100,000 times and compared the resulting correlation distribution against the observed value by computing a p-value at a significance level of 0.05.

To test (3) the significance of the groupings of climbing fiber branches based on Purkinje cell preferences, we performed a resampling test to simulate the condition in which climbing fibers established synaptic preferences onto random Purkinje cells among their observed targets. In each iteration of the resampling test we kept the identities of the Purkinje cells innervated by each climbing fiber branch fixed at observed values. Then, for each climbing fiber branch, we randomly sampled the number of synapses formed onto each of its targets from the observed synapse distribution (Figures 2c and 2d), with replacement. We only assessed ordering in our observed data for climbing fiber branches that formed a number of synapses onto one or more Purkinje cell targets that were in the top 10th percentile of the connectivity distribution. Thus, in order to make sure that the resampled distribution always contained at least one many-synapse connection, we repeated the resampling process for each climbing fiber branch until the above requirement was satisfied. We then ordered the climbing fiber branches with resampled connectivity using hierarchical clustering. We measured the tightness of the resulting clusters by computing the standard deviation of the distribution of distances between linked groups (i.e. the lengths of the vertical lines in Figure 7b). At each age we generated 10,000 resampled distributions in this way and compared the corresponding observed values by computing p-values at a significance level of 0.05.

### Scripts for Analysis

Scripts were written in Python 3.6, and can be found here (along with a list of package requirements): https://github.com/amwilson149/baby-andross.

### Dataset Hosting

Electron microscopy data is stored on Johns Hopkins University Applied Physics Laboratory’s BOSS (https://bossdb.org/project/PREwilson2019 [link will be updated].

