## Supplementary material for "The rich get richer: synaptic remodeling between climbing fibers and Purkinje cells in the developing cerebellum begins with positive feedback addition of synapses": All Supplemental Figures and Tables

### SUPPLEMENTAL INFORMATION

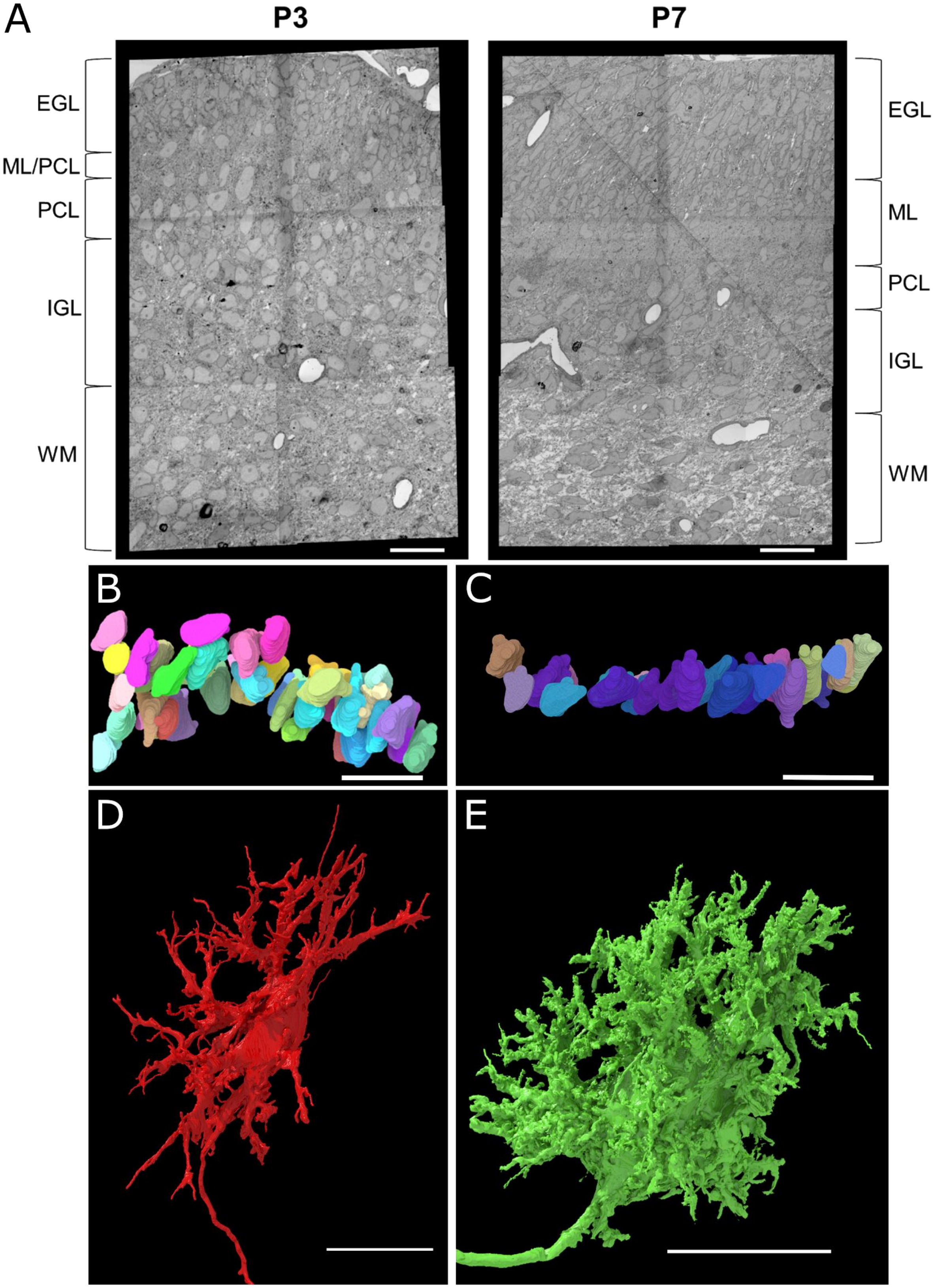
A
P3
P7
EGL
ML/PCL
PCL
IGL
WM
EGL
ML
PCL
IGL
WM
B
C
D
E

Figure S1. Locations of Purkinje Cells within the Layers of Cerebellar Cortex and their Detailed Morphologies

(a) Layers of the cerebellar cortex in example sections from the P3 (left) and P7 (right) datasets. The layers of the immature cerebellum vary in thickness between P3 and P7. In particular, the molecular layer grows as granule cells migrate to the internal granule layer, and Purkinje cells move from a multiple-layer to single-layer configuration during this time. Labels: EGL, external germinal layer; ML, molecular layer; PCL, Purkinje cell layer; IGL, internal granule layer; WM, white matter. These sections come from the regions of interest shown in Figure 1.

(b) Reconstructions of somas only for all Purkinje cells (48) in the P3 dataset, shown in the sagittal plane. At P3 the Purkinje cell layer was 2 to 3 cells deep.

(c) Reconstructions of somas only for all Purkinje cells (30) in the P7 dataset, shown in the sagittal plane. At P7 the Purkinje cell layer is a monolayer.

(d) Close-up, sagittal view of the fully reconstructed Purkinje cell at P3, showing the many dendrites present. The climbing fiber branches shown in Figures 1a and 1e formed their synapses mostly onto these dendrites.

(e) Close-up, sagittal view of the fully reconstructed Purkinje cell at P7, showing the many spines covering the soma and dendrites. The climbing fiber branches shown in Figures 1b and 1f formed their synapses mostly onto these spines.

Scale bars: (a) 20 µm; (b), (c) 30 µm; (d), (e) 15 µm.

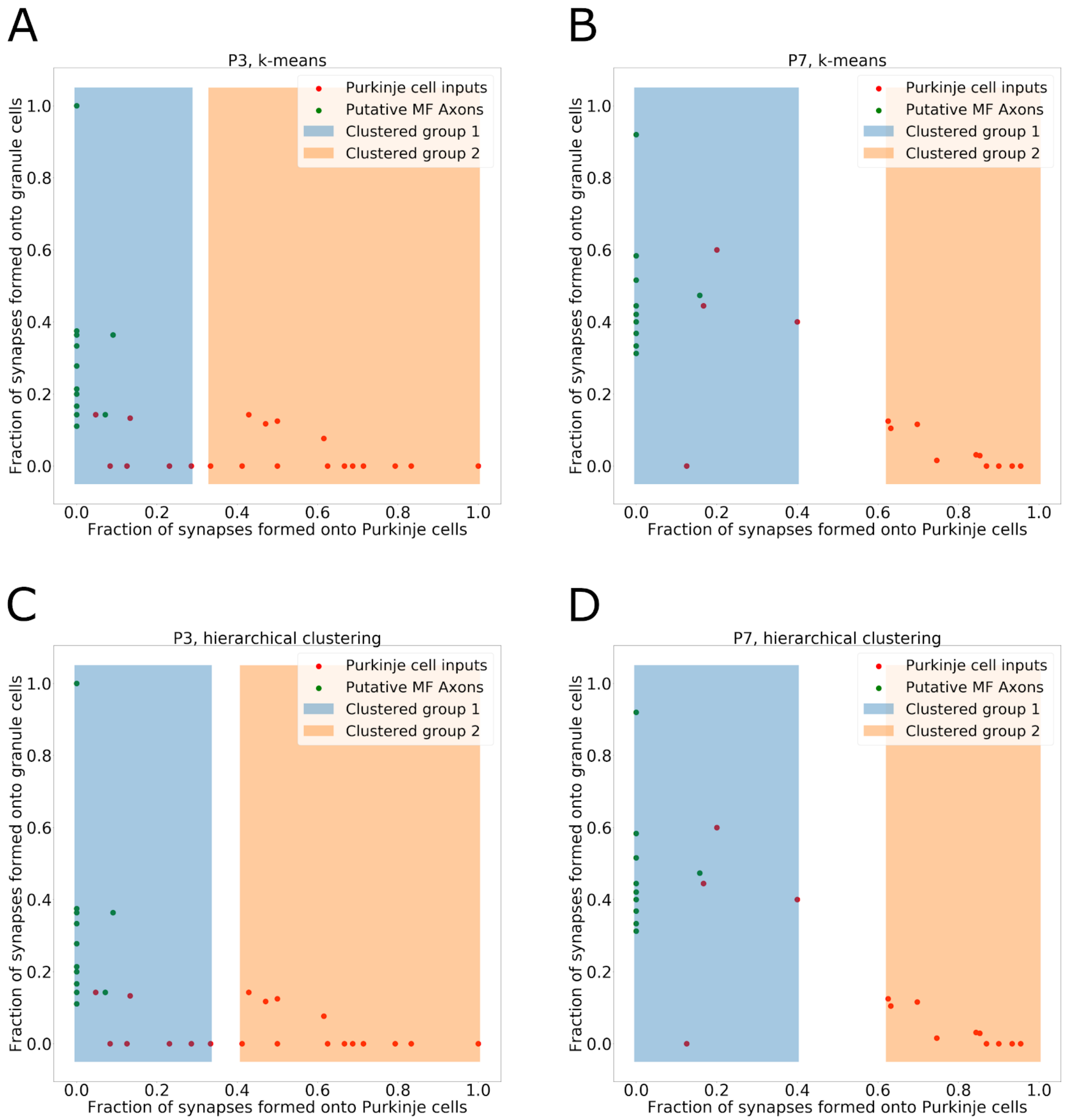


Figure S2. Classification of Non-Granule Excitatory Inputs to Fully Reconstructed Purkinje Cells Based on Synaptic Connectivity

(a) Results of k-means clustering performed on non-granule-excitatory axon branches (red data points) and putative mossy fiber axon branches (green points) at P3. Clustering was run to produce 2 groups. Shading indicates the boundary between the two groups, in the dimension corresponding to fraction of synapses formed onto Purkinje cells (group 1, blue shading; group 2, orange shading).

(b) Results of k-means clustering performed on non-granule-excitatory axon branches (red data points) and putative mossy fiber axon branches (green points) at P7. Clustering was run to produce 2 groups. Shading indicates the boundary between the two groups, in the dimension corresponding to fraction of synapses formed onto Purkinje cells (group 1, blue shading; group 2, orange shading).

(c) Results of hierarchical agglomerative clustering for non-granule excitatory axon branches and putative mossy fiber branches for P3. Clustering was used to generate two groups. Shading indicates the boundary between the two groups, in the dimension corresponding to fraction of synapses formed onto Purkinje cells (group 1, blue shading; group 2, orange shading).

(d) Results of hierarchical agglomerative clustering for non-granule excitatory axon branches and putative mossy fiber branches for P7. Clustering was used to generate two groups. Shading indicates the boundary between the two groups, in the dimension corresponding to fraction of synapses formed onto Purkinje cells (group 1, blue shading; group 2, orange shading).

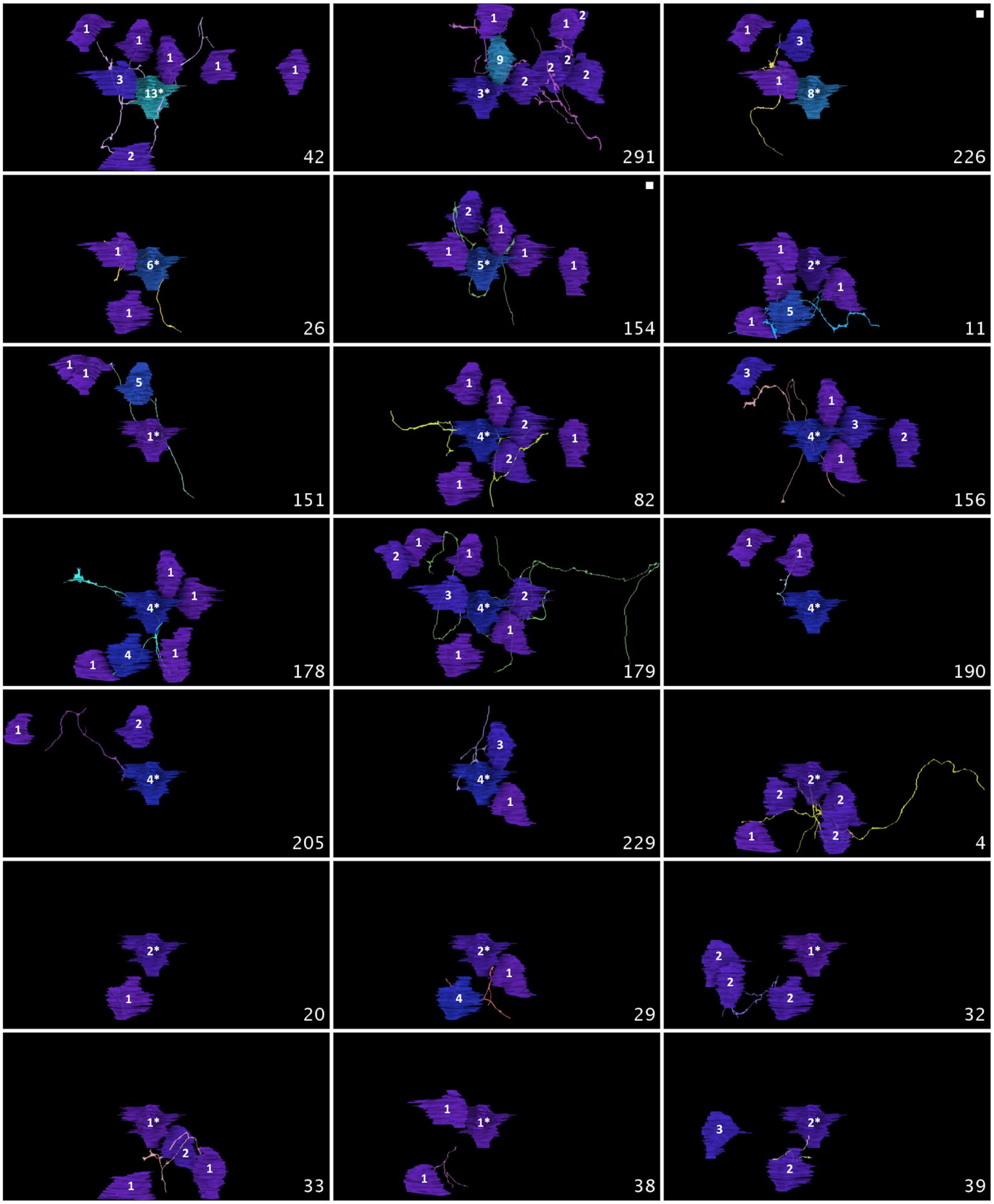
42
291
226
26
154
11
151
82
156
178
179
190
205
229
4
20
29
32
33
38
39

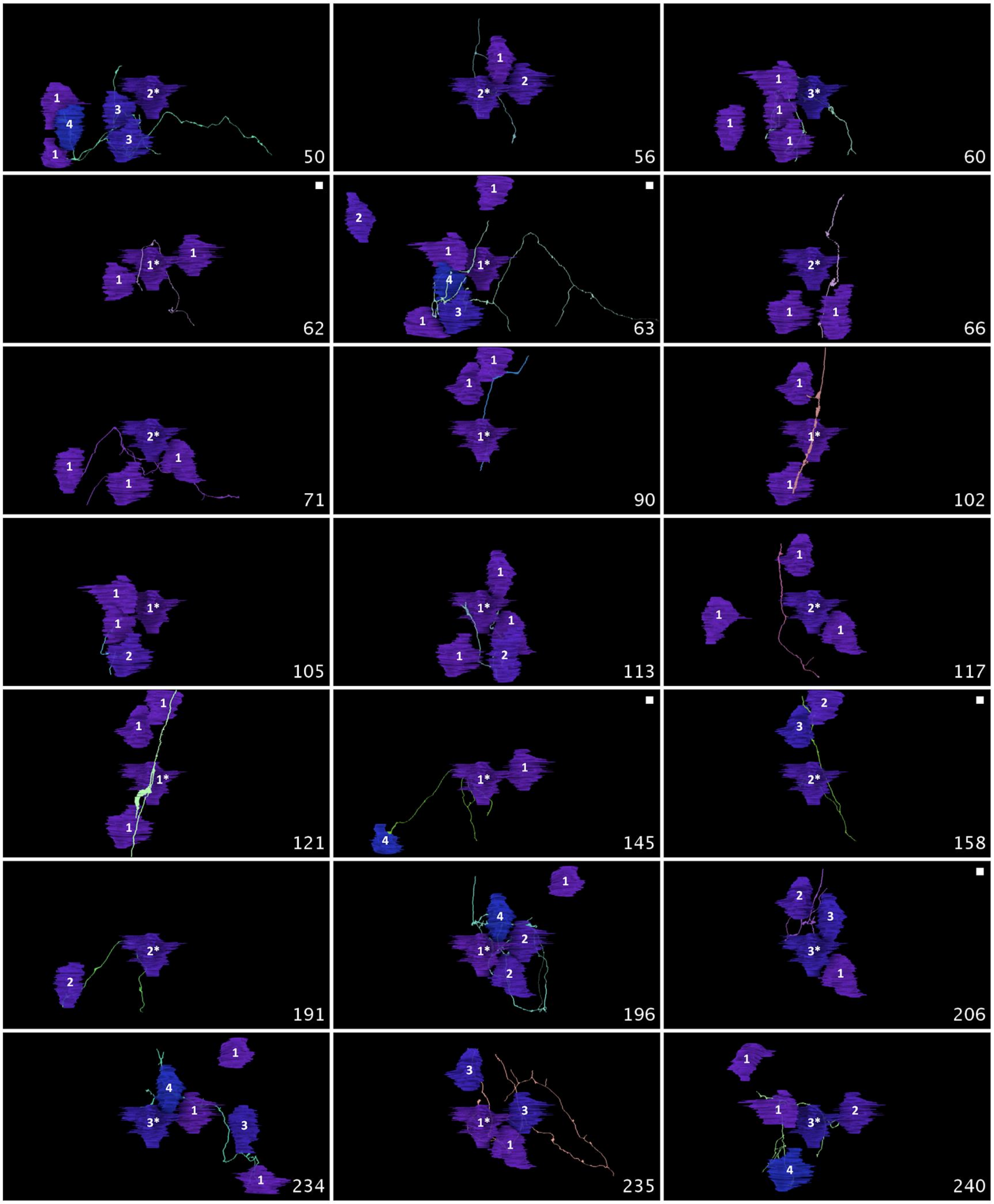
50
56
60
62
63
66
71
90
102
105
113
117
121
145
158
191
196
206
234
235
240

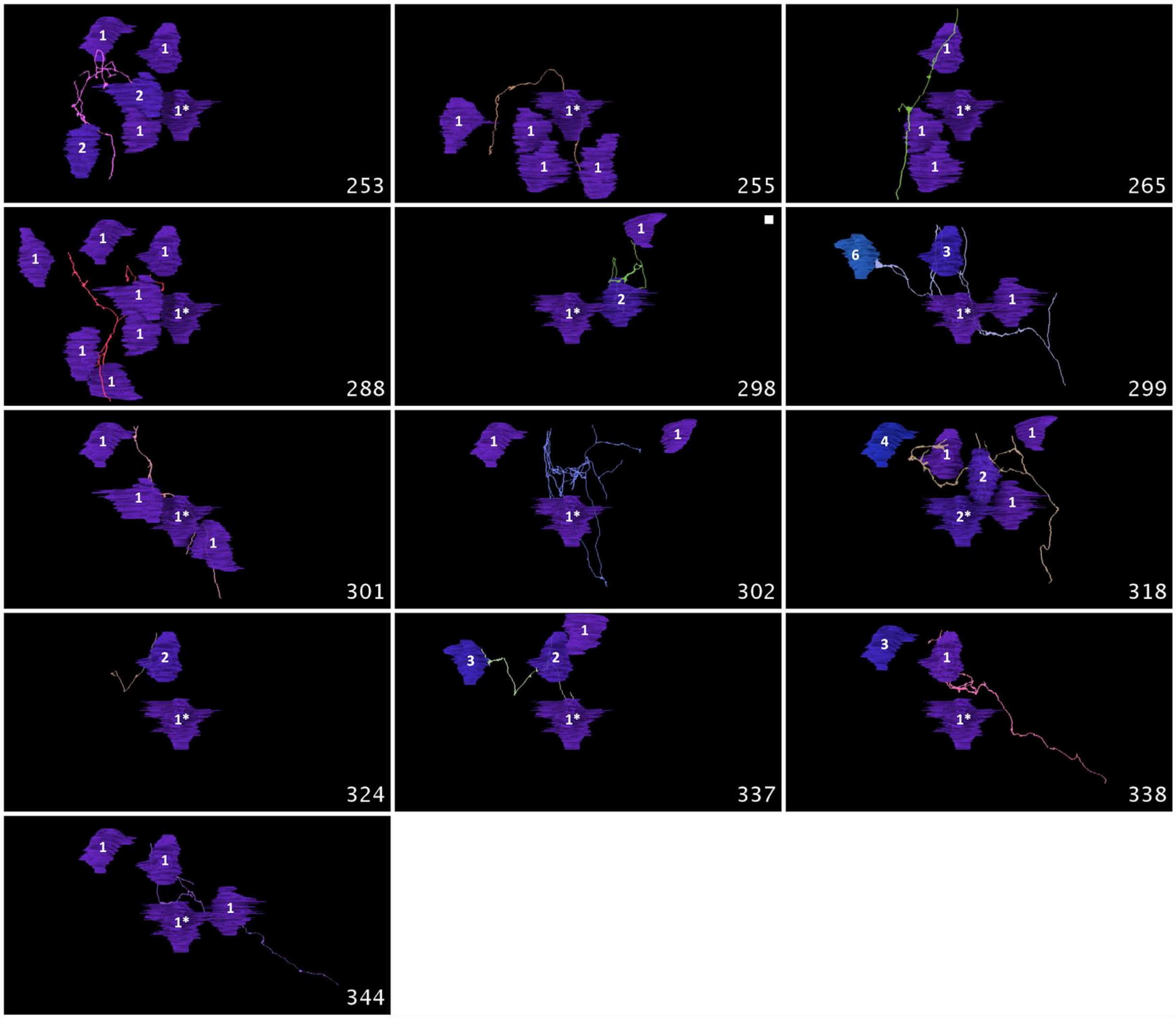


Figure S3. Connectivity for each of the 55 climbing fiber branches that innervate the fully reconstructed Purkinje cell at P3. Each branch is shown from the same view (looking from the pia toward the white matter) along with the Purkinje cells it innervates. The anteroposterior direction is horizontal. The number of synapses formed by a climbing fiber branch onto each Purkinje target is shown on the somata. Asterisks mark the fully reconstructed cell. Panels showing a climbing fiber branch with a full terminal arbor (i.e., one contained inside the P3 volume) have a white dot. The segment ID of the climbing fiber branch is shown in the bottom right corner of each image. A subset of these figures are shown in Figure 3a.

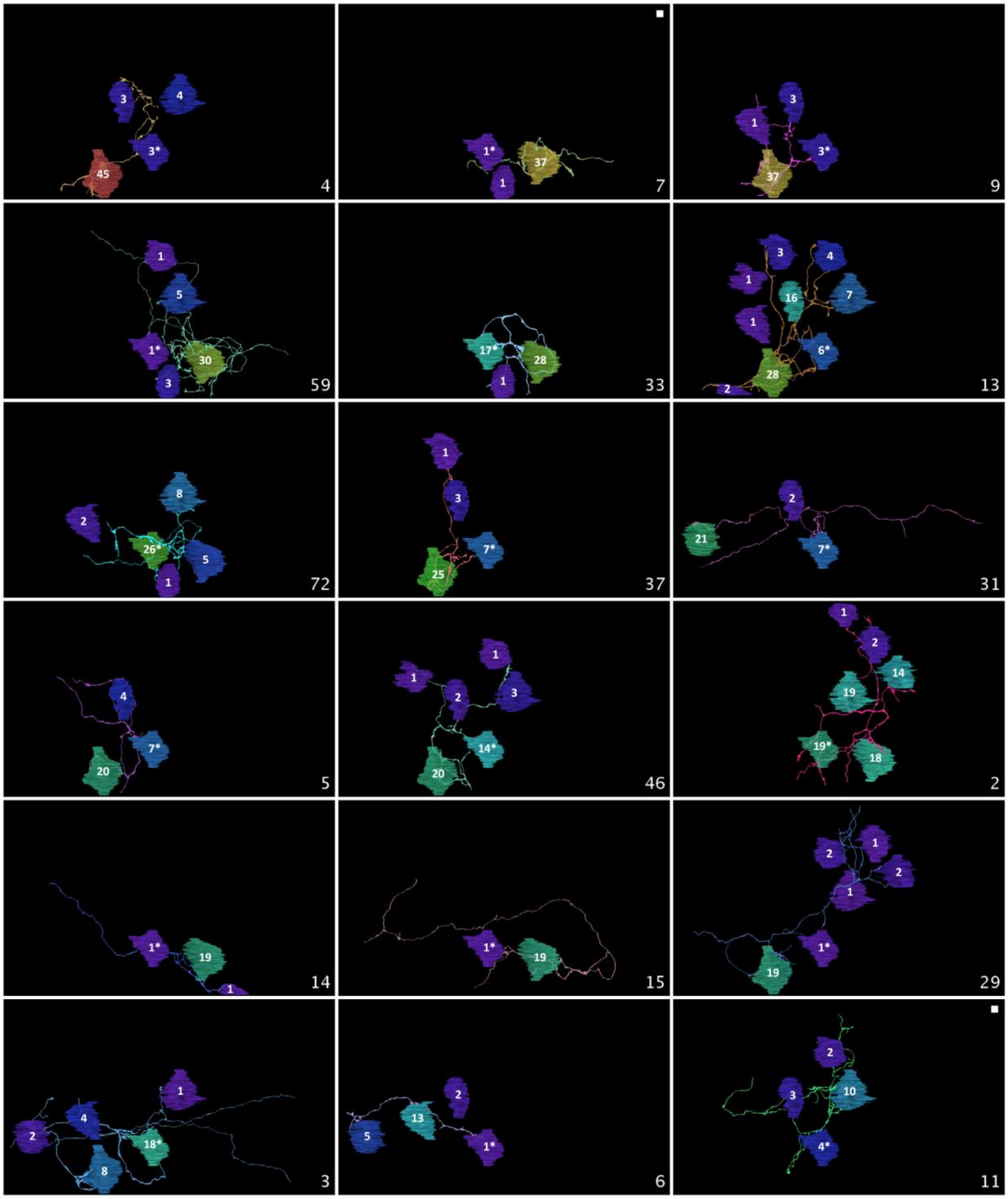
3
4
3*
45
4
1*
37
1
7
3
1
3*
37
9
1
5
1*
30
3
59
17*
28
1
33
3
4
1
16
7
1
6*
28
2
13
8
2
26*
5
1
72
1
3
7*
25
37
2
21
7*
31
4
7*
20
5
1
1
2
3
14*
20
46
1
2
14
19
19*
18
2
1*
19
1
14
1*
19
15
2
1
2
1
1*
19
29
1
4
2
18*
8
3
2
13
5
1*
6
2
3
10
4*
11

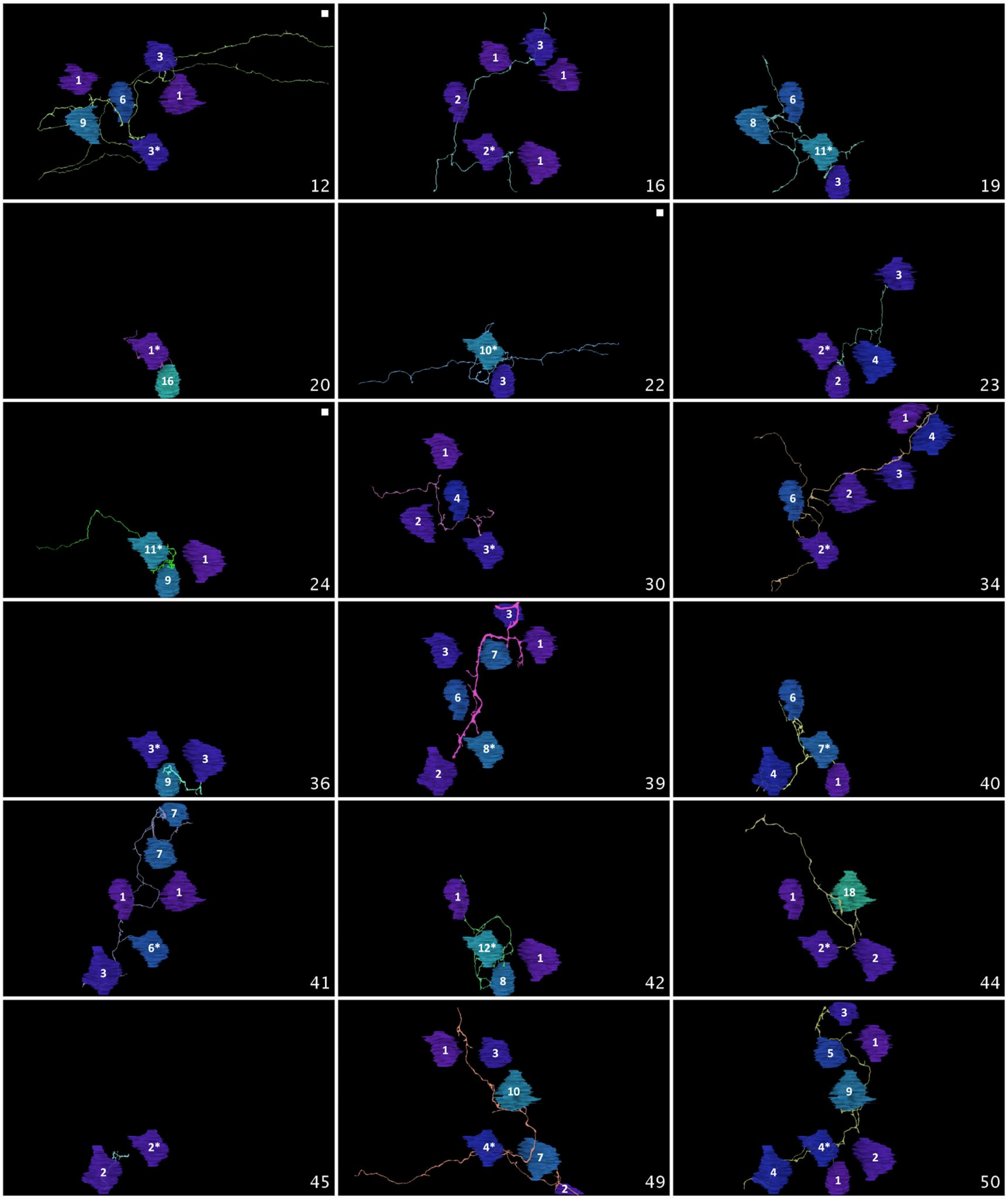
1 3 6 1 9 3* 12
1 3 1 2 2* 1 16
6 8 11* 3 19
1* 16 20
10* 3 22
3 2* 4 2 23
11* 1 9 24
1 4 2 3* 30
1 4 3 6 2 2* 34
3* 3 9 36
3 3 7 1 6 8* 2 39
6 7* 4 1 40
7 7 1 1 6* 3 41
1 12* 1 8 42
18 1 2* 2 44
2* 2 45
1 3 10 4* 7 2 49
3 1 5 9 4* 2 4 1 50

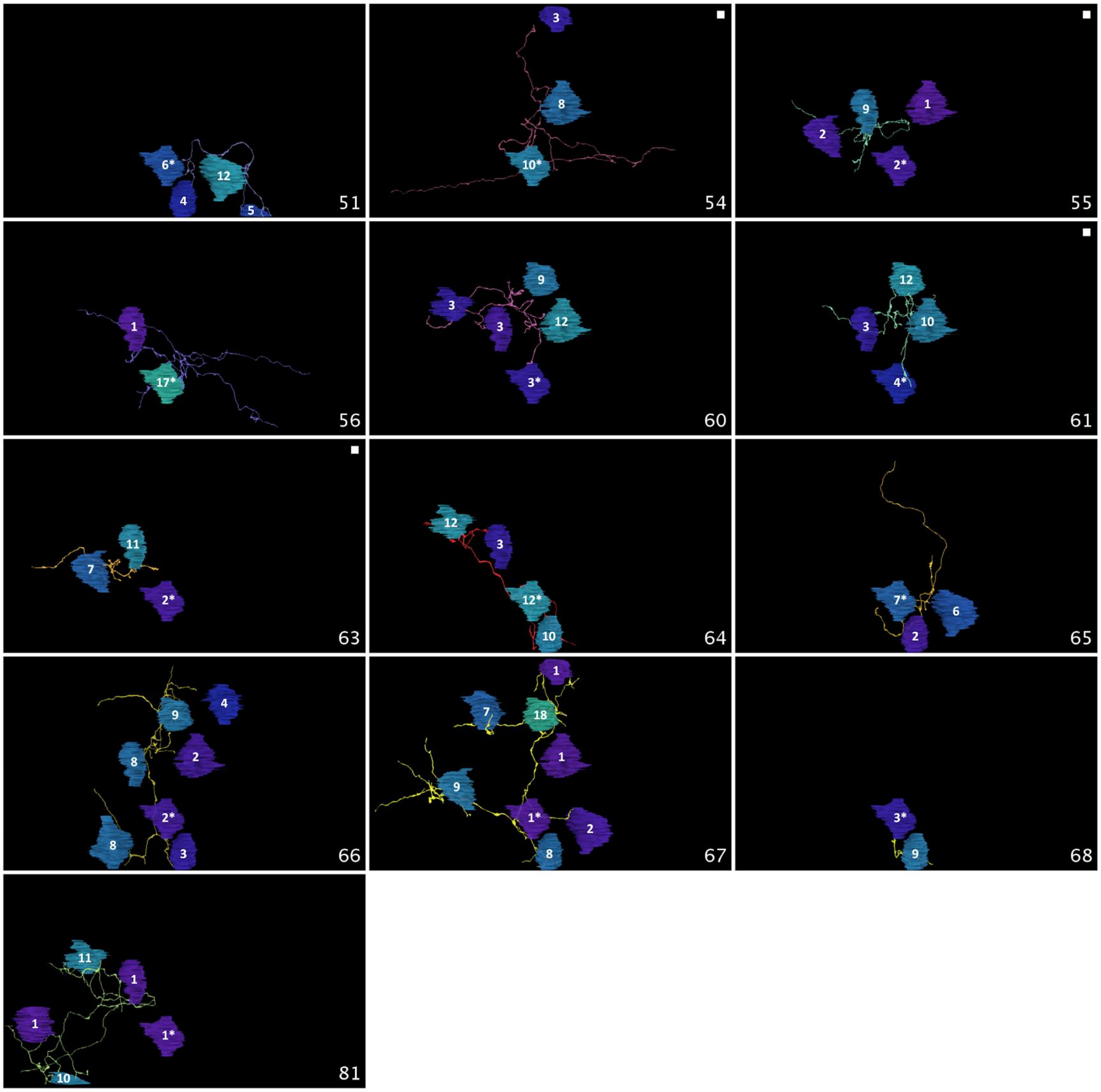


Figure S4. Connectivity for each of the 49 climbing fiber branches that innervate the fully reconstructed Purkinje cell at P7. Each branch is shown from the same view (looking from the pia toward the white matter) along with the Purkinje cells it innervates. The anteroposterior direction is horizontal. The number of synapses formed by a climbing fiber branch onto each Purkinje target is shown on the somata. Asterisks mark the fully reconstructed cell. Panels showing a climbing fiber with a full terminal arbor (i.e., one contained inside the P7 volume) have a white dot. The segment ID of the climbing fiber branch is shown in the bottom right corner of each image. A subset of these figures are shown in Figure 3a.

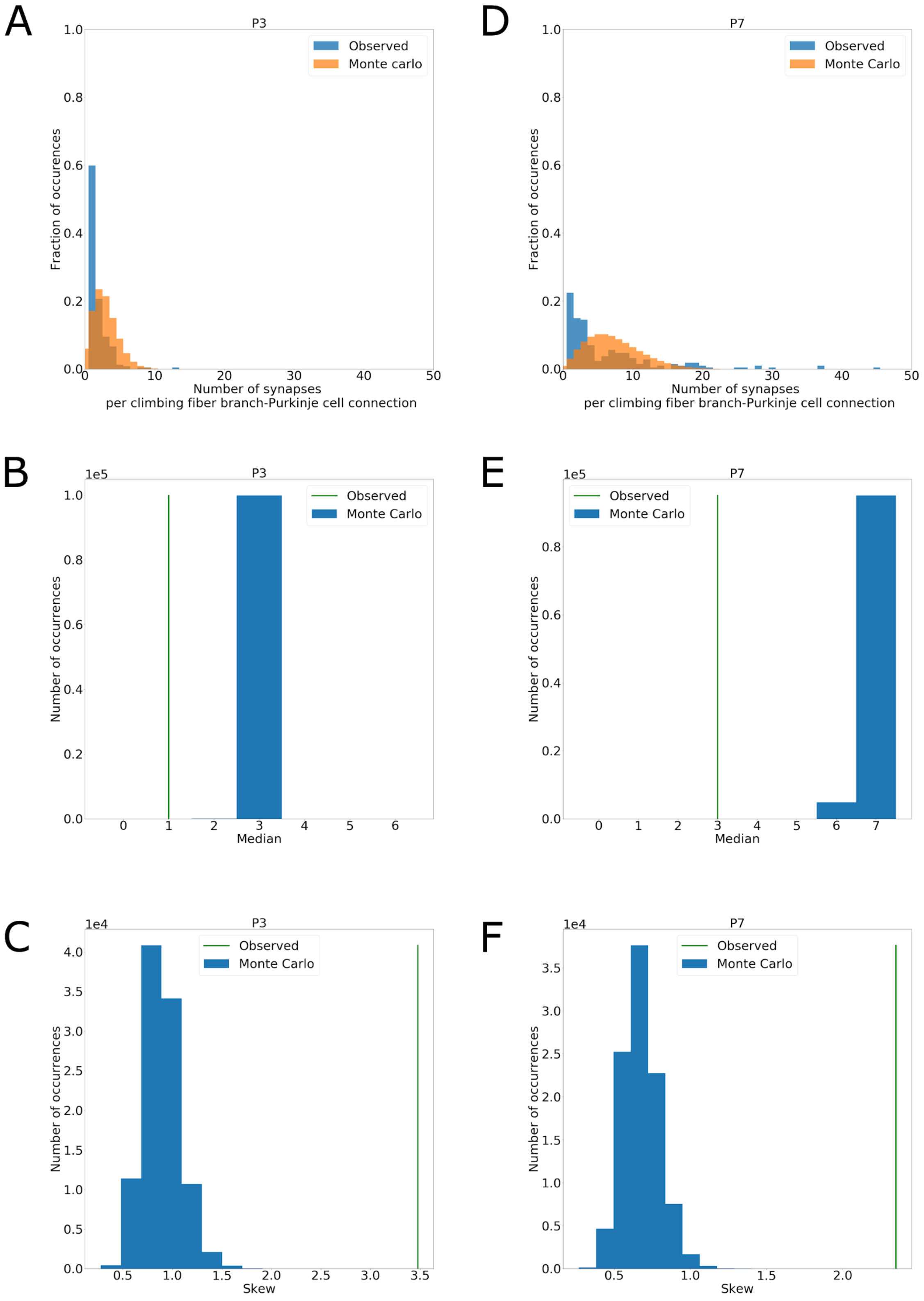
A
P3
Observed
Monte carlo
Fraction of occurrences
Number of synapses
per climbing fiber branch-Purkinje cell connection
D
P7
Observed
Monte Carlo
Fraction of occurrences
Number of synapses
per climbing fiber branch-Purkinje cell connection
B
1e5
P3
Observed
Monte Carlo
Number of occurrences
Median
E
1e5
P7
Observed
Monte Carlo
Number of occurrences
Median
C
1e4
P3
Observed
Monte Carlo
Number of occurrences
Skew
F
1e4
P7
Observed
Monte Carlo
Number of occurrences
Skew

Figure S5. Monte Carlo Test to Determine Whether Observed Synapse Distributions Could be the Result of Climbing Fiber Branches Innervating all Purkinje Cell Targets with Uniform Probability. See Methods and Figure 2.

(a-c) Results of a Monte Carlo simulation performed on P3 connectivity data.

(a) The observed connectivity histogram for all climbing fiber branch-Purkinje cell pairs analyzed (blue), compared with the Monte Carlo histogram (orange) generated when each climbing fiber branch was simulated as distributing its observed number of synapses onto its observed Purkinje cell targets with uniform probability. The Monte Carlo distribution shows the combined data from 100,000 simulations.

(b) The median of the observed connectivity distribution (green line) versus the medians of the Monte Carlo distributions (blue histogram).

(c) The skew of the observed (green line) versus Monte Carlo (blue histogram) distributions

(b-e) The same analysis for the P7 dataset.

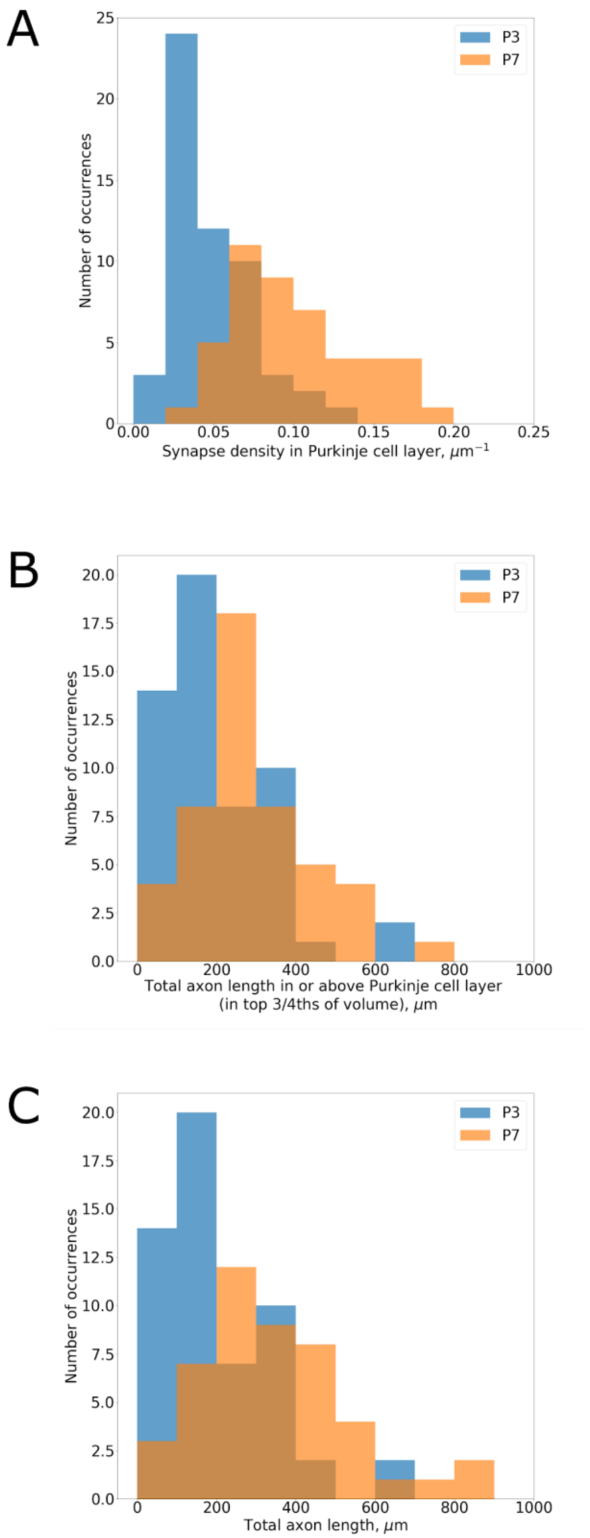


Figure S6. Lengths and Synapse Densities for Climbing Fiber Branches Reconstructed at P3 and P7.

(a) Histograms showing the density of synapses formed onto Purkinje cells within the Purkinje cell layer for climbing fiber branches reconstructed at P3 (blue) and at P7 (orange). The mean and standard deviation was 0.49 +/- 0.02 synapses/μm for the P3 distribution and 0.10 +/- 0.05 synapses/μm for the P7 distribution.

(b) Histograms showing the total axon branch length for each climbing fiber branch reconstructed at P3 (blue) and P7 (orange). These distributions are statistically different (Wilcoxon rank sum test, $p = 2 \times 10^{-5}$). The mean and standard deviation were 201 +/- 142 μm for the P3 distribution and 357 +/- 202 μm for the P7 distribution.

(c) Histograms showing the total climbing fiber branch length in the Purkinje cell layer for P3 (blue) and P7 (orange). These distributions are statistically different (Wilcoxon rank sum test, $p = 5.1\text{x}10^{-4}$). The mean and standard deviation was 197 +/- 138 μm for the P3 distribution and 292 +/- 146 μm for the P7 distribution.

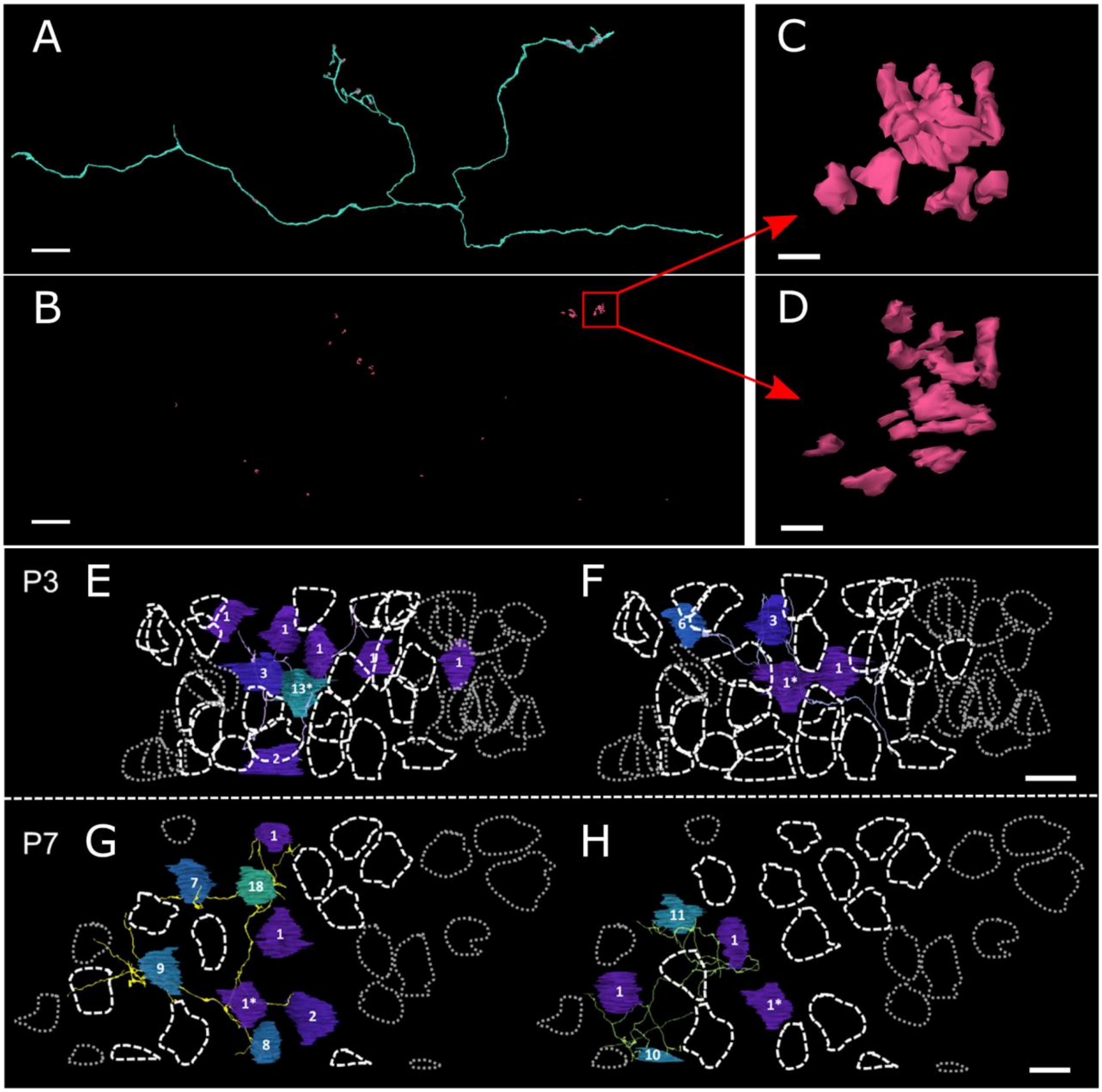


Figure S7. Patterns of Synaptic Innervation for Individual Climbing Fiber Branches.

(a) Three-dimensional rendering of a climbing fiber branch reconstructed at P7 (cyan, branch ID 31), along with all of its synapses (pink, ID 139), viewed from the sagittal plane.

(b) Three-dimensional rendering of only the synapses formed by the climbing fiber branch shown in (a). The red box indicates a region in which the climbing fiber forms a cluster of synapses.

(c) Close-up view of the cluster of synapses marked in (b), viewed from the same perspective as in (b). The climbing fiber branch forms multiple separate but closely apposed active zones.

(d) Close-up view of the cluster of synapses marked in (b), viewed along the dorsoventral axis (from the pia down to the white matter).

(e-h) Locations of Purkinje cells innervated by individual climbing fiber branches, relative to all other Purkinje cells in the volume (top panel, P3; bottom panel, P7). Dashed lines outline all Purkinje cells that are innervated by the climbing fiber inputs to the fully reconstructed Purkinje cell. Dotted lines outline Purkinje cells in the volume that are not innervated by those climbing fiber branches. Asterisks identify the fully reconstructed Purkinje cell. This figure is related to Figures 3a, S3, and S4.

(e) One of the reconstructed climbing fiber branches in the P3 image volume (light purple axon; branch ID 42) and the Purkinje cells it innervates (colored-in Purkinje cell somata also showing the number of synapses between them). The dendrites of the Purkinje cells are not shown. The terminal arbor of this axon innervates a number of Purkinje cells but they are not all contiguous with each other. For example the rightmost and lowest Purkinje cells are surrounded by Purkinje cells that are not innervated by this axon.

(f) Another example of the sporadic innervation of Purkinje cells by an individual P3 climbing fiber branch (periwinkle, ID 299).

(g, h) Examples of innervation patterns for two climbing fiber branches (yellow, ID 67, and green, ID 81) and their Purkinje cell targets.

Scale bars: (a), (b) 7.5 μm; (c), (d) 500 nm; (e)-(h) 15 μm.

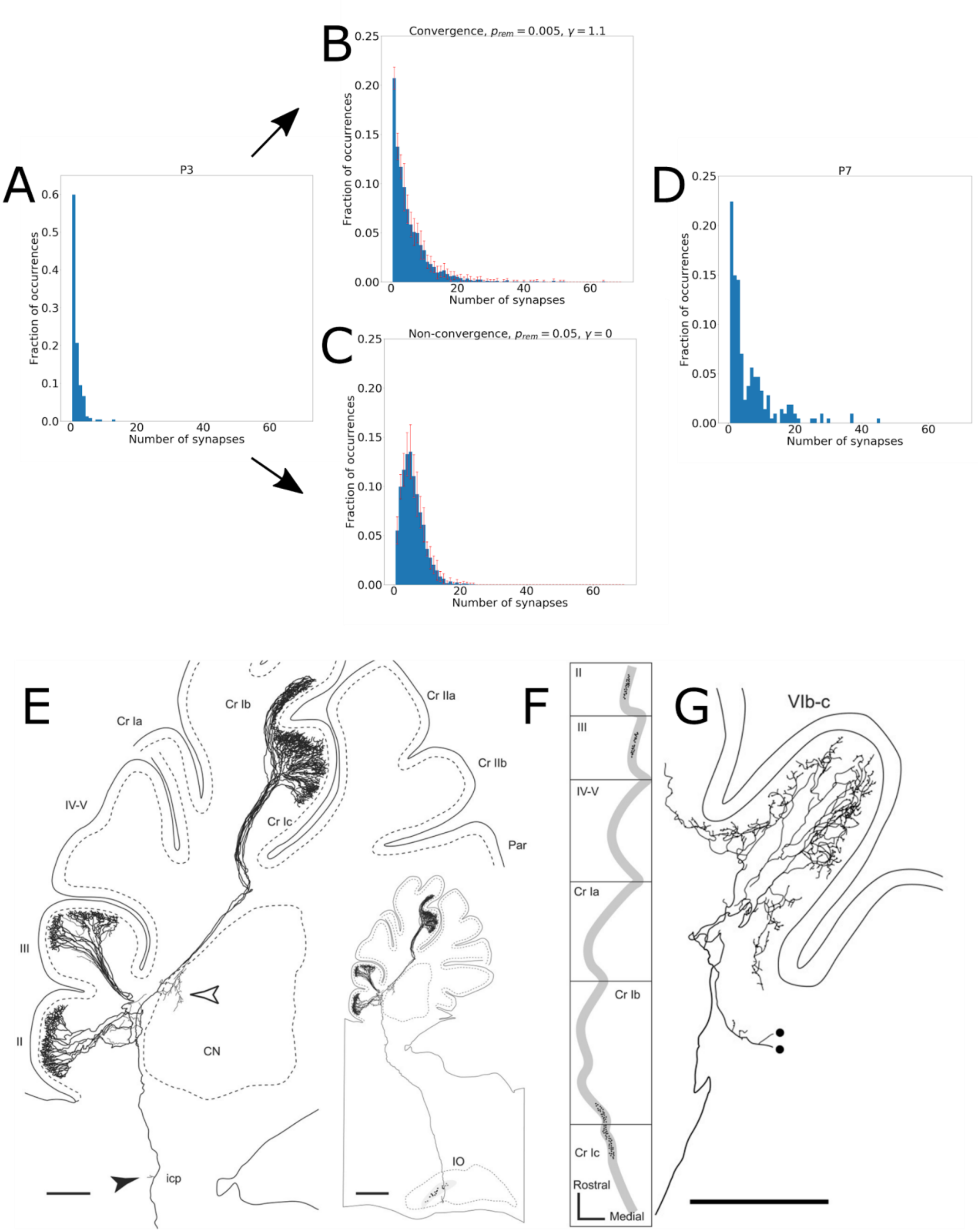


Figure S8. Additional Details Relating to Simulations.

(a-d) Examples of connectivity distributions produced by the time-evolution simulation with parameter values that led to convergence versus with parameter values that did not

(a) Histogram of the initial connectivity distribution; i.e. the observed P3 distribution from Figure 2c.

(b) Mean +/- standard deviation of the connectivity distribution produced by the simulation using parameters that lead to convergence ($p_{rem}$ = 0.005, $\gamma$ = 1.1), at a time step where convergence is at its best (955 time steps; see text). This distribution was produced based on 11 simulation runs.

(c) Mean +/- standard deviation of the connectivity distribution produced by the simulation using parameters that never led to convergence ($p_{rem}$ = 0.05, $\gamma$ = 0), run to the same number of time steps as in (b) for comparison (i.e. 955). This distribution was produced based on 11 simulation runs.

(d) Histogram of the target connectivity distribution (i.e. the observed P7 distribution), for comparison with (b) and (c).

(e-g) Reproductions of Figures 1 ((e), (e) inset, and (f)) and Figure 6a ((g)) from (Sugihara, 2005). The densities of terminal arbors formed by single climbing fiber axons reconstructed in this work (see e.g. middle panel) were used to estimate the number of climbing fiber axons innervating the fully reconstructed P7 Purkinje cell.

(e) A single labeled climbing fiber axon.

(f) A radial view of the climbing fiber axon in (e). For this axon 24, 27, and 62 terminal arbors were reported in (Sugihara, 2005) in lobules II, III and crus Ib-c, respectively.

(g) A single climbing fiber axon that extended 24 terminal arbors into lobule VIb-c (Figure 6a in Sugihara, 2005).

Scale bars: (e), (g) 200 μm; (e) inset, (f) 500 μm.

Table S1. Numbers of synapses formed by non-granule-cell excitatory axon branches and mossy fiber branches onto all identified target types in the P3 dataset.

| P3 | | | | |
|---|---|---|---|---|
| **Axon segment ID** | **Population** | **N syns, Purkinje cells (PCs)** | **N syns, granule cells (GCs)** | **N syns, confirmed non-PC, non-GC cells** |
| 59 | PC input | 2 | 0 | 0 |
| 60 | PC input | 7 | 0 | 0 |
| 176 | PC input | 2 | 2 | 6 |
| 288 | PC input | 8 | 2 | 5 |
| 302 | PC input | 3 | 1 | 3 |
| 305 | PC input | 3 | 0 | 1 |
| 346 | PC input | 3 | 0 | 1 |
| 357 | PC input | 1 | 0 | 0 |
| 358 | PC input | 1 | 3 | 2 |
| 365 | PC input | 1 | 0 | 2 |
| 26 | PC input | 8 | 0 | 4 |
| 42 | PC input | 23 | 0 | 3 |
| 62 | PC input | 3 | 0 | 0 |
| 71 | PC input | 5 | 0 | 0 |
| 82 | PC input | 11 | 0 | 1 |
| 105 | PC input | 5 | 0 | 1 |
| 145 | PC input | 6 | 0 | 0 |
| 151 | PC input | 8 | 1 | 1 |
| 255 | PC input | 5 | 0 | 2 |
| 265 | PC input | 4 | 1 | 2 |
| 324 | Putative MF | 0 | 6 | 0 |
| 325 | Putative MF | 0 | 7 | 7 |
| 327 | Putative MF | 0 | 2 | 1 |
| 322 | Putative MF | 0 | 8 | 1 |
| 355 | Putative MF | 0 | 3 | 2 |
| 402 | Putative MF | 0 | 3 | 0 |
| 522 | Putative MF | 0 | 18 | 15 |
| 521 | Putative MF | 1 | 2 | 1 |
| 511 | Putative MF | 1 | 4 | 2 |
| 461 | Putative MF | 0 | 2 | 2 |
| 321 | Putative MF | 0 | 5 | 3 |
| 328 | Putative MF | 0 | 1 | 0 |

Table S2. Numbers of synapses formed by non-granule-cell excitatory axon branches and mossy fiber branches onto all identified target types in the P7 dataset.

| P7 | | | | |
|---|---|---|---|---|
| **Axon segment ID** | **Population** | **N syns, Purkinje cells (PCs)** | **N syns, granule cells (GCs)** | **N syns, confirmed non-PC, non-GC** |
| 40 | PC input | 18 | 0 | 1 |
| 32 | PC input | 12 | 2 | 4 |
| 55 | PC input | 14 | 0 | 0 |
| 49 | PC input | 27 | 1 | 4 |
| 61 | PC input | 29 | 1 | 1 |
| 24 | PC input | 21 | 0 | 0 |
| 15 | PC input | 20 | 0 | 1 |
| 21 | PC input | 30 | 5 | 4 |
| 10 | PC input | 5 | 1 | 1 |
| 67 | PC input | 47 | 1 | 4 |
| 25 | PC input | 1 | 3 | 0 |
| 26 | PC input | 2 | 2 | 0 |
| 35 | PC input | 1 | 0 | 3 |
| 53 | PC input | 3 | 8 | 3 |
| 71 | Putative MF | 3 | 9 | 4 |
| 72 | Putative MF | 0 | 7 | 7 |
| 73 | Putative MF | 0 | 2 | 1 |
| 75 | Putative MF | 0 | 6 | 8 |
| 591 | Putative MF | 0 | 8 | 3 |
| 311 | Putative MF | 0 | 23 | 1 |
| 361 | Putative MF | 0 | 5 | 7 |
| 371 | Putative MF | 0 | 8 | 3 |
| 381 | Putative MF | 0 | 16 | 1 |
| 431 | Putative MF | 0 | 14 | 2 |
